# Deep-Learning Super-Resolution Microscopy Reveals Nanometer-Scale Intracellular Dynamics at the Millisecond Temporal Resolution

**DOI:** 10.1101/2021.10.08.463746

**Authors:** Rong Chen, Xiao Tang, Zeyu Shen, Yusheng Shen, Tiantian Li, Ji Wang, Binbin Cui, Yusong Guo, Shengwang Du, Shuhuai Yao

**Affiliations:** Department of Chemical and Biological Engineering, Hong Kong University of Science and Technology, Hong Kong, China; Division of Life Science, The Hong Kong University of Science and Technology, Hong Kong, China; Department of Mechanical and Aerospace Engineering, Hong Kong University of Science and Technology, Hong Kong, China; Department of Physics, The Hong Kong University of Science and Technology, Hong Kong, China; Department of Physics, The University of Texas at Dallas, Richardson, Texas 75080, USA

**Author notes:** Correspondence and requests for materials should be addressed to Y.G., S.D., or to S.Y.

## Abstract

Single-molecule localization microscopy (SMLM) can be used to resolve subcellular structures and achieve a tenfold improvement in spatial resolution compared to that obtained by conventional fluorescence microscopy. However, the separation of single-molecule fluorescence events in thousands of frames dramatically increases the image acquisition time and phototoxicity, impeding the observation of instantaneous intracellular dynamics. Based on deep learning networks, we develop a single-frame super-resolution microscopy (SFSRM) approach that reconstructs a super-resolution image from a single frame of a diffraction-limited image to support live-cell super-resolution imaging at a ∼20 nm spatial resolution and a temporal resolution of up to 10 ms over thousands of time points. We demonstrate that our SFSRM method enables the visualization of the dynamics of vesicle transport at a millisecond temporal resolution in the dense and vibrant microtubule network in live cells. Moreover, the well-trained network model can be used with different live-cell imaging systems, such as confocal and light-sheet microscopes, making super-resolution microscopy accessible to nonexperts.

## Introduction

Live-cell fluorescence imaging, requiring both low phototoxic illumination and a high imaging speed, is usually performed with a wide-field (WF) fluorescence microscope ^1^. The spatial resolution of a conventional fluorescence microscope is limited by the lightwave diffraction effect and thus unable to resolve subcellular structures smaller than 200 nm. In the past two decades, various types of super-resolution microscopy surpassing the diffraction limit have been developed. For example, structured illumination microscopy (SIM) ^2^ can be used for live-cell imaging with low invasiveness; however, it only improves the spatial resolution of images by a factor of up to 2 and requires multiple frames to construct a single super-resolution (SR) image. Stimulated emission depletion (STED) microscopy ^3^ can achieve an ∼50 nm resolution using highly intense light pulses, but point-to-point scanning makes STED too slow for live-cell imaging. Single-molecule localization microscopy (SMLM), including photoactivated localization microscopy (PALM) ^4^ and stochastic optical reconstruction microscopy (STORM) ^5^, further enhances the spatial resolution by a factor of ten (∼20 nm) but typically requires more than thousands of frames with separated single-molecule fluorescence events to reconstruct one SR image; hence, in rare cases, SMLM has been applied to live cells at a second-scale temporal resolution ^6, 7^. To perform time-resolved and noninvasive super-resolution imaging, numerous advanced labeling strategies ^8-10^, optical imaging systems ^11, 12^, and image reconstruction methods ^13-15^ have been explored in recent decades. Nonetheless, inherent tradeoffs among spatial and temporal resolutions, the achievable signal intensity and cytotoxicity must be made due to the physical boundaries of optical systems ^16^.

The rapid development of artificial intelligence has led to many traditional hardware limits being surpassed. Various deep learning networks have displayed excellent performance in the single-image super-resolution (SISR) task ^17-19^ which usually transform a single low-resolution (LR) photograph to a high-resolution (HR) photograph. The focus of the SISR task for realistic photographs is to enhance texture and improve visual quality ^20, 21^. In contrast, super-resolution tasks for microscopic images involve ultrastructure recovery from diffraction-limited images with high accuracy. Recently, popular neural networks in computer vision have been modified to enhance the microscopic image resolution. However, limited resolution improvements have been achieved, for instance, from low magnification to high magnification ^22, 23^, confocal to STED, and total internal reflection fluorescence (TIRF) or WF to SIM ^23-26^. At even higher resolutions, such as those achieved by PALM and STORM, although deep-learning methods have been reported to accelerate the localization process of STORM reconstruction ^27^ and reduce the number of frames of single-molecule images required for PALM reconstruction ^28^, multiple frames with single-molecule fluorescence events are still required for the reconstruction of an SR image. Therefore, the fundamental problems of multiframe super-resolution imaging, such as the long acquisition time and photobleaching-induced phototoxicity in localization microscopy, still hinder its application in the imaging of live-cell dynamics.

In this work, we first explore the possibility of using a neural network to directly transform a diffraction-limited image to an SR image at a single-molecule resolution. By applying an enhanced super-resolution generative adversarial network (ESRGAN) ^21^, multicomponent loss function, and prior information regulation, we develop a super-resolution network (SRN) that can resolve a single diffraction-limited frame to a SR image with up to a 10-fold resolution improvement. Then, we investigate the challenges of implementing this SRN for real-time live-cell observation involving images with an ultralow signal-to-noise ratio (SNR). By deploying a signal-enhancement network (SEN) in advance to progressively optimize the image SNR and resolution, we successfully decouple the input image SNR with the final reconstruction quality, thus allowing for high-speed live-cell imaging without sacrificing the spatial resolution. Taken together, we propose a single-frame super-resolution microscopy (SFSRM) approach that allows us to reveal time-resolved intracellular events in live cells at a 20 nm spatial resolution and a 10 ms temporal resolution for thousands of time points. As a reference, our SFSRM is applied to visualize millisecond cargo transport dynamics in the dense microtubule (MT) network inside living cells, which provides valuable insights into live, subtle, and complex vesicle behaviors that were previously unresolvable. Moreover, we demonstrate that the well-trained SFSRM networks can be used in various imaging systems without further training, making super-resolution imaging possible for labs lacking training datasets.

## Results

### SFSRM based on a joint-optimization-enhanced deep learning network

The performance of a deep-learning network is mainly influenced by three components: the network architecture, the training loss function, and the training data. Since a three-layer neural network was first utilized in enhancing image resolution ^17^, the network architecture has been expanded ^18, 19, 29^; notably, a deep and complex structure is proven to facilitate the description of the relationship between LR and HR images, hence producing superior results ^20^. However, as the network depth increases, training becomes particularly challenging, and the result may degrade due to overfitting or training failure. To solve this problem, ESRGAN utilizes residual-in-residual dense blocks to effectively train deep networks ^21^. A residual channel attention network (RCAN)^30^ proposes the “channel attention” mechanism to guide the network to focus on high-frequency details. Recently, a detailed-fidelity attention network (DeFiAN) ^31^ combines the channel-attention mechanism ^30^ with feature filtering ^31^ to reconstruct high-frequency details and to maintain high-accuracy. In our extreme super-resolution task, we selected three abovementioned networks as candidates and tested their performance in reconstructing the ground-truth (GT) image from a 10-fold degraded LR image. The reconstruction results in **Fig. 1b(I))** indicate that ESRGAN successfully resolves two polylines that are only approximately 20 nm apart; however, the other two networks fail to reconstruct the subtle structures. We repeated the experiment on 30 images and quantitatively assessed the network reconstruction fidelity based on the multiscale structural similarity (MS-SSIM) between the network output and the GT image. The statistical results indicate that ESRGAN also achieves higher fidelity than the other two networks (**Fig. 1b(I)**, box-and-whisker plot). Therefore, we adopted ESRGAN as the basic network of the developed SFSRM method (**Fig. 1a**).

**Figure 1:**
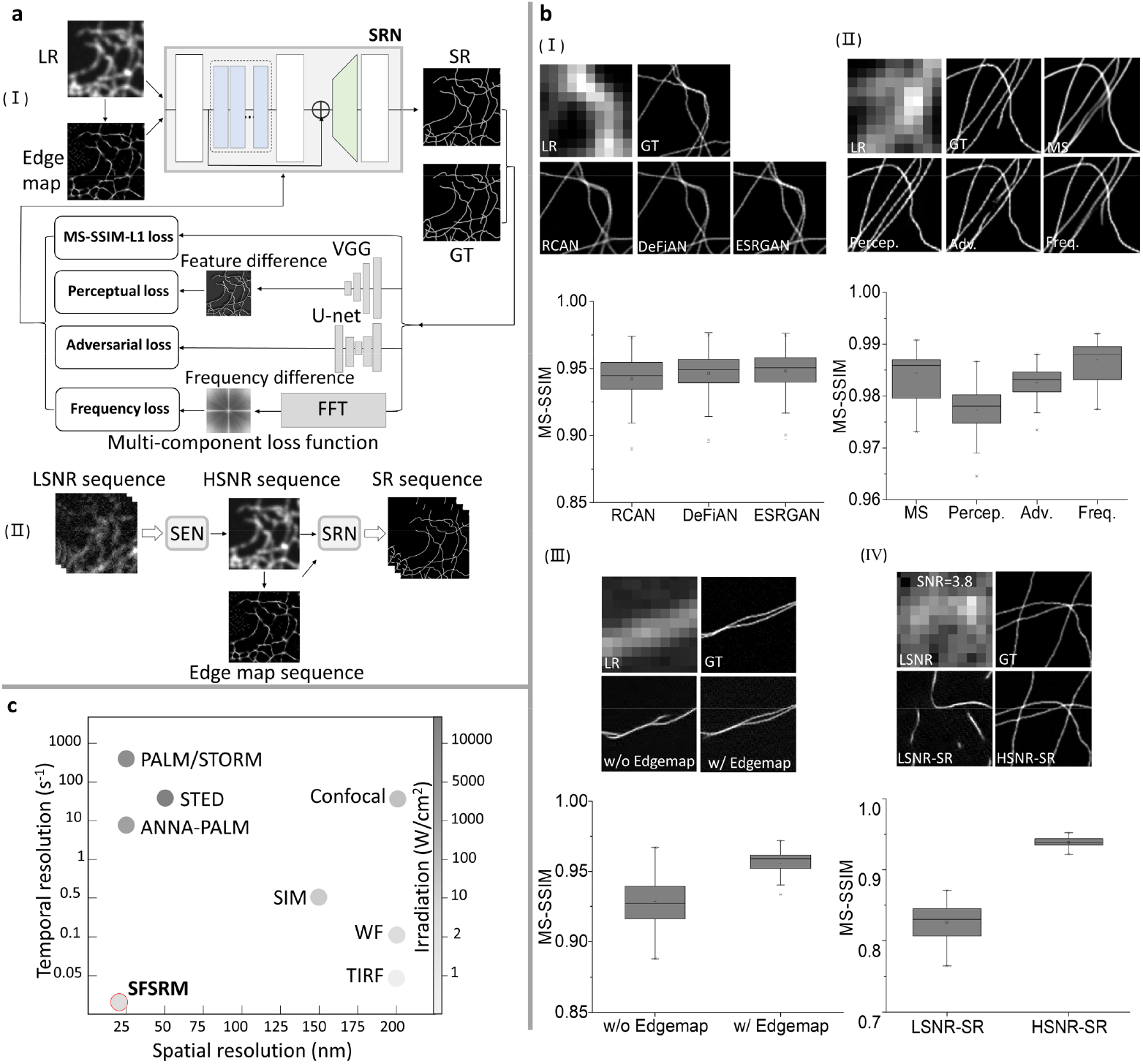
Overview of SFSRM. **(a)** The networks of SFSRM. (I) The super-resolution network (SRN) architecture. The SRN is trained using simulated low-resolution (LR) image and ground-truth (GT) image pairs or experimental wide-field (WF) and GT image pairs obtained stochastic optical reconstruction microscope (STORM). The LR/WF image is preprocessed by a sub-pixel edge-detector to extract the edge map, and both of them are then fed as inputs to the network. A multi-component loss function including: (i) MS-SSIM-L1 loss which measures the pixel-wise difference between the SR and GT image; (ii) Perceptual loss which measures the difference between feature maps of SR and GT images extracted by the VGG network; (iii) Adversarial loss returned by the discriminator which distinguishes the GT image from the SR image; (iv) Frequency loss which compares the frequency spectrum difference of the SR and GT image in a specified frequency region, is adopted to train the network. (II) Workflow of SFSRM. To implement SFSRM for high frame-rate live-cell imaging, a low-SNR (LSNR) image sequence acquired from a living cell first goes through the signal-enhancement network (SEN) to improve signal intensity (or SNR), and the intermediate result (HSNR image sequence) is further processed to extract the edge map and input to the SRN for reconstruction of a super-resolution (SR) image. **(b)** Ablation study of different components in SFSRM. The network reconstruction fidelity is measured by multi-scale structure similarity (MS-SSIM) index between the reconstructed images and the GT image. The error bar represents reconstruction experiments repeated on 30 images. (I) Comparison of three network architectures including Residual Channel Attention Network (RCAN), Detail-Fidelity Attention Network (DeFiAN), and Enhanced Super Resolution Generative Adversarial Network (ESRGAN). (II) Comparison of four loss function components. MS, Percep., Adv. and Freq. are the aberrations for MS-SSIM-L1 loss, perceptual loss, adversarial loss, and frequency loss respectively. (III) Comparison the reconstruction results with (w/) and without (w/o) the edge map. (IV) Comparison the network reconstructions from the LSNR image (SNR=3.8) and the HSNR image. **(c)** Spatiotemporal and irradiation intensity map of the available microscopy including photo-activated localization microscopy (PALM), STORM, stimulated emission depletion (STED) microscopy, artificial neural network accelerated PALM (ANNA-PALM), Confocal, structured illumination microscopy (SIM), WF, total internal reflection fluorescence (TIRF) microscopy, and SFSRM, most of which suffer from the tradeoff between spatial and temporal resolution. In contrast, SFSRM achieves a spatial resolution of 20 nm and a temporal resolution of 10 ms.

We note that ESRGAN yields abundant high-frequency details due to its network architecture and the consideration of perceptual loss. Perceptual loss have been widely used in enhancing the resolution of realistic photograph but not been applied in super-resolving microscopic image because although visually pleasing textures are obtained, fuzzy regions and artifacts may be created ^20^. In microscopic image restoration, in pursuit of high accuracy, the most commonly used loss functions, including mean absolute error (MAE), mean square error (MSE), and structural similarity (SSIM) losses, focus on pixelwise differences between the network output and the GT image. Although these loss functions can provide high peak signal-to-noise ratio or structure similarity (SSIM) index values, they are often limited by oversmoothing and lose high-frequency details ^20^. Thus, to date, most super-resolution networks applied to microscopic images can only achieve limited resolution improvements ^23-26^. Here, we introduce perceptual loss for the microscopic image restoration for the first time to achieve high-frequency detail reconstruction. To guarantee the accuracy of reconstruction details, we developed a multicomponent loss function that combines pixel-oriented loss (MS-SSIM-L1 loss) and perceptual loss to improve the fidelity of details; additionally, adversarial loss and frequency loss are combined to penalize blur in images and suppress high-frequency artifacts. The performance of different loss components is shown in **Fig. 1b(II)**. Unsurprisingly, using the pixel-oriented MS-SSIM-L1 loss function alone results in oversmoothed filaments, which are more apparent at cross points. Adding perceptual loss aids in the reconstruction of fine lines that are visually similar to GTs; however, the accuracy of the structure remains similar to that based on MS-SSIM-L1 loss. Further adding adversarial loss aids in the reconstruction of some correct structures but also induces artifacts that are corrected by incorporating frequency loss. We note that the MS-SSIM-L1 loss function yields the highest MS-SSIM index in the MS-SSIM comparison (**Fig. 1b(II)**, chart). However, this result does not necessarily suggest that the MS-SSIM-L1 loss provides the highest accuracy since errors are difficult to detect in blurry images. In contrast, perceptual loss and adversarial loss provide sharp reconstructed structures, and slight mismatch between these results and the GT results can be detected. Thus, the MS-SSIM index decreases after adding perceptual loss and adversarial loss. As frequency loss further improves the reconstruction accuracy by removing the artifacts induced by adversarial loss, the MS-SSIM index increases again. Overall, by considering each loss component, the well-trained network is able to restore fine structures with a 10× resolution improvement and achieves an MS-SSIM index of 0.98 with respect to the GT in noise-free conditions, manifesting the capability of the network to transform a single diffraction-limited image to an SR image with a 10-fold resolution increase.

Unfortunately, the reconstruction result quickly degrades if the image is disturbed by noise (**Fig. 1b(III)**, without the edge map), which is reasonable since single-image super-resolution restoration is already an ill-posed problem and noise will add further complexity to this task. Determining how to improve the reconstruction accuracy of noisy images remains a critical task. Here, we innovatively integrate the prior information (edge map) from an LR image into the network to aid in reconstruction. Although edge priors have been considered in realistic photograph restoration^32, 33^, edge detection operators that are well suited for realistic photographs cannot be directly applied to microscopic images since the diffraction effect is not considered. As shown in **Supplementary Fig. 1a**, the edges extracted from the microscopic LR image by these operators fail to indicate the high-resolution structures in the GT image. Instead, we extracted a subpixel edge map from a microscopic image based on the radial symmetry of the fluorophore (**Supplementary Fig. 1b**) which has been utilized in super-resolution microscopy methods ^34-36^. Inspired by Gustafsson et al., who analyzed the temporal cumulates in the radiality maps of a sequence of images to reconstruct one SR image ^34^, we computed the edge map from a single LR image (**Supplementary Fig. 1c**) and used it as an additional input to the network. The result in **Fig. 1b(III)** indicates that the edge map successfully guides the network to restore fine features from the noisy LR image and enhance accuracy (plot in **Fig. 1b(III)**).

We further validated our conclusions based on experimental images of microtubules, and the result exhibited good agreement with the simulation data (**Supplementary Fig. 2**). We then explored whether the SRN can maintain superiority when processing signals with different densities. As shown in Supplementary **Fig. 3**, a series of images with increasing signal density (see the Methods for the signal density quantification) was used to test the performance of the SRN. The relationship between the reconstruction error based on the normalized root-mean-square error (NRMSE) and the signal density is plotted in **Supplementary Fig. 3b**. As expected, the NRMSE increases as the signal density increases. Nonetheless, the NRMSE of the SR reconstruction remains at an acceptable error level, even for the highest signal density (NRMSE = 0.07, signal ratio = 0.6). Based on a visual comparison between the reconstructed image and the GT image, we empirically set NRMSE = 0.07 as an error threshold for follow-up experiments. Moreover, the SRN consistently achieves a resolution comparable to that of the GT image, regardless of the signal density (**Supplementary Fig. 3c**). The excellent performance of the SRN prompts us to ask whether the network can be applied to images with low SNRs. As shown in **Supplementary Fig. 4a** (LSNR and LSNR-SR), we assessed the SRN performance based on a sequence of images with increasing SNR and quantitatively analyzed the relationship between the input image SNR and the reconstruction error. **Supplementary Fig. 4b** (LSNR-SR) shows that the reconstruction error increases as the input image SNR decreases. When the input image SNR<14, the reconstruction error exceeds 0.07, indicating that the network does have an SNR requirement for LR images. This prerequisite for SNR poses a challenge for applications in live-cell imaging, in which the SNR is generally very low due to the short exposure time and low illuminance. To address this challenge, we employed a signal-enhancing network (SEN) to improve the image SNR in advance (**Fig. 1c**). For SEN training, we acquired low-SNR (LSNR) and high-SNR (HSNR) image pairs of fixed cells at different illumination intensities and from different microscopes, including epifluorescence, TIRF, highly inclined and laminated optical sheet (HILO), and confocal microscopes, to allow the network to learn the statistical pattern of the noise and effectively extract the structural information from LSNR images. As shown in **Supplementary Fig. 4b** (HSNR-SR), the well-trained SEN is able to extend the minimum requirement to SNR of 7, thus allowing for high-speed and low-irradiation live-cell imaging (see Supplementary Note 1 for more details about SNR estimation and measurement). In addition, the SEN enhances the robustness of the network for different imaging systems. As demonstrated in **Supplementary Fig. 5**, where the same cell is imaged by a Zeiss sp8 confocal microscope to obtain a confocal image and a Zeiss Elyra 7 microscope in HILO mode to obtain a WF image and STORM mode to obtain a GT image, the network successfully restores the SR images from the two LR images without degradation due to the low SNR of the confocal image or high background of the WF image; therefore, the capability of this network to be effectively applied in various systems is demonstrated.

As a consequence of joint optimization from different perspectives, our SFSRM can not only resolve ultrastructures with single-molecule resolution benefiting from SR microscopy, but also track live-cell dynamics at high imaging speed and low photobleaching by virtue of the WF microscope (**Fig. 1c**). Therefore, this approach is a powerful tool combining the merits of the super-resolution microscopy and real-time live-cell imaging.

### SFSRM reconstructs a super-resolution image from a single diffraction-limited image

Before applying SFSRM for live-cell imaging, we validated its performance based on experimental images of fixed cells. **Figs. 2a-b** show the original WF image of MTs stained with Alexa Fluor 647 and the reconstruction result of SFSRM. We compared the performance of SFSRM and a representative deep-learning-based super-resolution method called ANNA-PALM ^28^, which involves two trained networks: one trained with a single WF image, denoted as ANNAPALM (WF), and another trained with a WF image and a sparse-PALM (SP) image reconstructed from 300 frames of single-molecule images, called ANNAPALM (WF+SP). The reconstruction results of the two networks are shown in **Figs. 2c-d**. The differences between the network reconstruction results and the GT image (**Fig. 2e**) are illustrated (**Figs. 2f-h**). From this comparison (**Figs. 2a-e**), we found that when the MTs are sparse and isolated, all networks can perfectly reconstruct the MTs; however, when the signal is dense and MTs cross or are close to each other, as demonstrated in the two zoomed-in regions (**Figs. 2i-j**), the MTs tended to be merged, blurred, or lost in the ANNAPALM (WF) reconstruction. Although the sparse-PALM image improves the structure patterns in general, the multiframe acquisition of sparse-PALM images leads to reduced imaging speed and additional photobleaching, thus hampering live-cell imaging. In contrast, our SFSRM can reconstruct the MTs from a WF image without losing filaments in dense regions or merging filaments when they are close to each other, demonstrating performance similar to that of ANNAPALM (WF+SP). For example, **Fig. 2k** shows the signal distribution along the lines indicated in **Fig. 2i**; the five densely distributed filaments are successfully reconstructed by SFSRM, whereas three of them are lost in the ANNAPALM (WF) and ANNAPALM (WF+SP) reconstruction results. As depicted by the intensity profiles for the lines in **Fig. 2j**, two MTs only 120 nm apart are merged in the ANNAPALM (WF) reconstruction result, separated but shifted in the ANNAPALM (WF+SP) reconstruction result, and correctly resolved in the SFSRM reconstruction result. We quantitatively assess the reconstruction results of the networks based on three metrics: the MS-SSIM, NRMSE, and resolution measured by image decorrelation analysis ^37^. SFSRM exhibits drastically better performance than ANNAPALM (WF), with improved fidelity (0.892 vs. 0.867), lower reconstruction error (0.0651 vs. 0.0842), and an approximately twofold-higher resolution (43 nm vs. 78 nm) (**Fig. 2m**).

**Figure 2:**
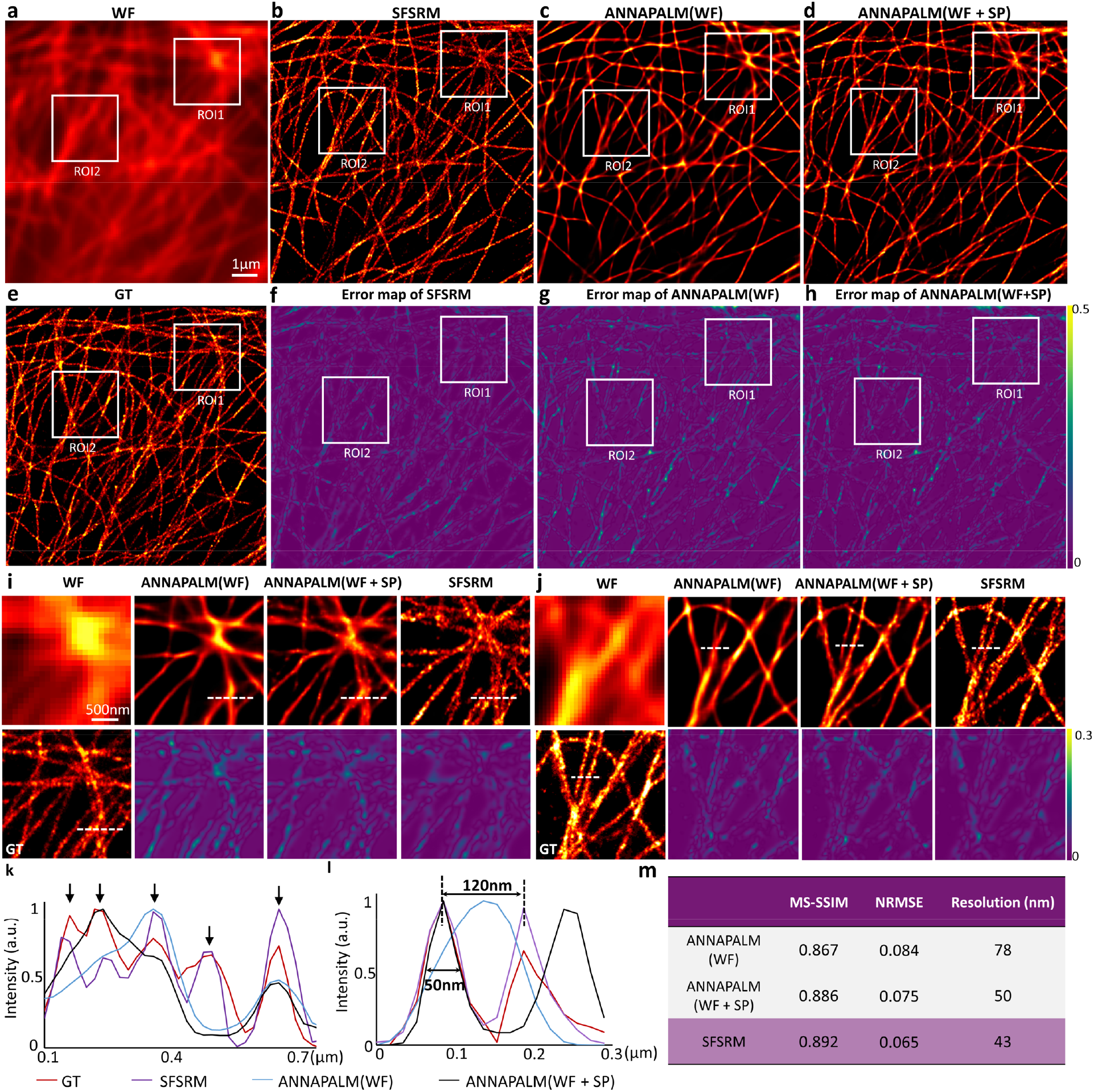
SFSRM reconstruction from a single WF image of immunostained microtubules. **(a)** WF image. **(b)** SFSRM reconstruction from the WF image. **(c)** ANNA-PALM reconstruction from the WF image, noted as ANNA-PALM (WF). **(d)** ANNA-PALM reconstruction from the WF image and a sparse PALM image obtained from 300 frames of single-molecule images, noted as ANNA-PALM (WF+SP). **(e)** STORM image obtained from 20,000 frames of single-molecule images, noted as GT. **(f-h)** Difference images of the SFSRM reconstruction, ANNA-PALM (WF) reconstruction, and ANNA-PALM(WF+SP) reconstruction with respect to the GT image. **(i, j)** Zoom-in regions of interest ROI1 and ROI2 in **a-h. (k, l)** Intensity profiles along the lines indicated by the white dashed lines in **i** and **j**. Black arrows in **k** indicate the five microtubules close to each other are successfully reconstructed by SFSRM while some of them are missed in ANNA-PALM (WF) and ANNA-PALM (WF+SP) reconstruction. The width of the microtubules (∼50 nm) and the spacing of adjacent microtubules (∼120 nm) are indicated in **j. (m)** Quantitative assessment of three reconstruction results. Reconstruction fidelity is assessed by multi-scale structure similarity (MS-SSIM). Reconstruction error is quantified via normalized root-mean-square error (NRMSE). Reconstruction resolution is evaluated by decorrelation analysis.

In addition to using our own data, we applied the trained SFSRM approach to a publicly available simulated dataset and an experimental dataset from the EPFL SMLM challenge website ^38^. The reconstruction results of SFSRM are comparable to those of Deep-STORM ^27^, which requires several hundred densely distributed single-molecule images (**Supplementary Fig. 6**), demonstrating the high robustness of the proposed approach for various signal densities and different sets of simulated and experimental data.

### SFSRM reveals the millisecond dynamics of cargo trafficking in live cells

Intracellular transport plays an essential role in maintaining cellular functions ^39^. Many cellular processes rely on the transport system to deliver proteins or organelles to a specific functional location. External cargos such as viruses and nanoparticles also utilize the transport system to deliver their genomes or drugs to specific compartments for function ^40^. Considering that the cytoplasm of eukaryotic cells is highly crowded and dynamic, how cargo is delivered across the cytoplasm to specific positions remains largely unclear. Previous studies have reported that MTs form a complex network throughout cells and serve as highways to deliver cargo between the perinuclear region and the cell periphery in rapid and directed motions involving motor proteins ^41, 42^. Recently, facilitated by single-particle tracking techniques ^43-45^, mounting dynamic behaviors during the cargo transport process, e.g., back-and-forth movement, rotation, pause, and switching direction, have been discovered, suggesting that rapidly directed motions are frequently interrupted. Some in vitro studies have suggested that the intersections of the microtubules are likely to interfere with cargo transport and form tethering points for cargo ^46, 47^. Single-particle tracking combined with confocal microscopy or STORM microscopy has also been employed to investigate vesicle behavior at microtubule intersections in live cells ^43, 44, 48, 49^. However, confocal microscopy fails to provide a high-resolution MT map, and STORM requires the sequential imaging of vesicles and MTs. In addition, to register the vesicle trajectory along MTs in STORM images, MT dynamics are stabilized by paclitaxel and nocodazole during live-cell imaging ^48-50^. The lack of real-time high-resolution MT imaging has impeded the further exploration of cargo-MT interactions. Hence, the underlying mechanism of the complicated dynamics of vesicular trafficking along MTs remains largely unknown. Our SFSRM approach will facilitate the simultaneous visualization of cargo and MT dynamics with unprecedented spatiotemporal resolution in live cells, thereby providing comprehensive insight into cargo transport involving MTs.

Here, as a demonstration, we investigate the dynamics of the endocytic trafficking of epidermal growth factor (EGF) protein in live cells. Considering the different spectra involved in fixed-cell and live-cell labeling, we first validated the robustness of our network for different spectra (**Supplementary Fig. 7**). To visualize EGF protein and MT in living cells, networks were pretrained with MT and EGF images separately (**Supplementary Fig. 8**). For live-cell imaging, cells were transfected with plasmids encoding mEmerald-tagged MT-associated fusion protein ensconsin (3XmEmerald-ensconsin) ^51^ and then incubated with Qdot655-labeled EGF protein to monitor EGF endocytosis. SFSRM imaging at a high speed (100 Hz) and a low illumination intensity (15 W/cm^2^) for 5000 time points clearly reveals the EGF and MT dynamics, from which we identified the novel dynamics of MT that were unexplored in previous studies (**Supplementary Video 1**, part I). With the unprecedented resolution improvement (**Fig. 3a**, WF vs. SR), the tangled MT network is clearly resolved, and MTs with various morphologies, such as bending, crossing, and bundles, can be observed (**Figure 3a**; SR). Furthermore, MT deformation dynamics, such as MT bending (**Fig. 3b, Supplementary Video 1**, part II), growth and shrink instability (**Fig. 3d, Supplementary Video 1**, part II), are recorded at an ultrafast imaging speed, which allows us to capture high-frequency fluctuations, including the time-varying MT bending (**Fig. 3c**; 100 Hz) and the random-walk MT growth trajectory (**Fig. 3e**; 100 Hz). These results suggest that the intracellular dynamics at the millisecond scale may be greatly underestimated at a low sampling frequency (**Fig. 3c and 3e**; 2 Hz). Moreover, we reveal for the first time the intracellular MT transverse vibration at a high frequency in the approximately 100 nm range (**Fig. 3f, Supplementary Video 1**, part III), which is undetectable in the WF image. The statistical analysis of the transverse MT displacement in 10 ms intervals (**Fig. 3f**, histogram) shows that the typical displacement is approximately 30 nm, which is far larger than the system drift (less than 10 nm in 50 seconds; **Supplementary Fig. 9**), thus confirming our observation regarding true MT vibration. The rapid and random vibration of MTs could cause bundle instability (**Fig. 3g**), or the local MT network morphology change(**Fig. 3h**).The observed MT vibration, which may result from fluctuations in the embedded viscoelastic actin cytoskeleton ^52^, the dynamic load of traveling motor proteins ^53^ (**Supplementary Fig. 10**), or the hydrolysis of guanosine triphosphate^54^, is involved in multiple cellular functions, such as organizing and maintaining cell shape ^55^, promoting cilia movement ^56^, and mediating cargo transport; therefore, this function is of high research interest. However, due to the previous resolution limitations, in vivo MT fluctuations could only be inferred from the motion of bound motor proteins ^57^. Currently, the direct visualization of MT vibration may promote the comprehension of intracellular force fluctuations^58^ and provide evidence for different models of microtubule vibration ^59, 60^.

**Figure 3:**
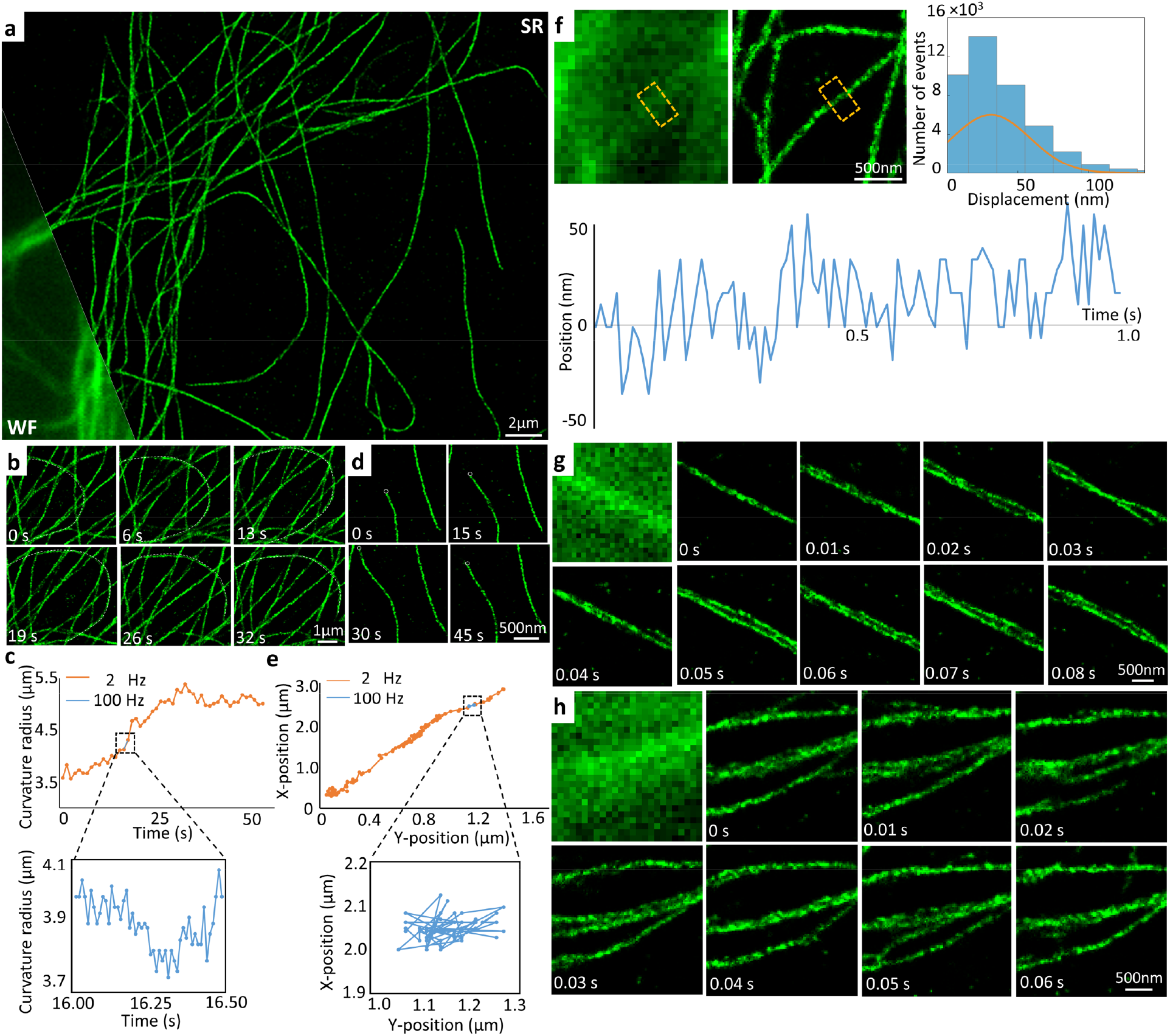
Real-time SFSRM imaging reveals microtubule dynamics. **(a)** Representative image of microtubules (green) from cells expressing mEmerald-ensconsin. Bottom left: a fraction of the corresponding WF image. **(b, c)** Microtubule bending dynamics and the curvature radius recorded at 2 Hz and 100 Hz. **(d, e)** Microtubule tip growth and shrink dynamics and tip displacement recorded at 2 Hz and 100 Hz. **(f)** Microtubule transverse fluctuation. Top left: comparison of the raw live-cell image and SR reconstruction of microtubules. Top right: histogram of microtubule transverse displacement at a 10 ms interval. Bottom: microtubule fluctuation over time recorded at 100 Hz. **(g)** Microtubule bundle instability caused by inconsonant vibrations of MTs. **(h)** Local microtubule topology change in a short time.

Next, we focus on the influence of the observed MT dynamics on cargo transport. The EGF endocytosis process was recorded with dual-channel SFSRM (**Fig. 4a, Supplementary Video 2**, part I). High-spatiotemporal-resolution videometry (50 nm resolution for MTs, 20 nm resolution for EGF, and 100 Hz imaging speed) clearly reveals the vesicle transport details, from which we noticed that slight fluctuations of a single MT does not interrupt the directed transport of vesicles, but the motions of the vesicles along the fluctuating MTs are significantly more dynamic than expected. **Fig. 4b(I-III)** illustrates three examples of vesicle transport dynamics: (I) moving back and forth along an MT, (II) rotating around an MT, and (III) colliding with other vesicles and then changing direction (**Supplementary Video 2**, part II). These subtle and fast random walks are undetectable at low spatial (**Fig. 4c, I**) and temporal (**Fig. 4c, II**) resolutions (**Supplementary Video 2**, part II), which implies that vesicle movement is scale dependent. At the millisecond scale, thermal diffusion is dominant (**Fig. 4c, III**; 100 Hz, α = 0.25), and at the second scale, directed transport dominates (**Fig. 4c, III**; 100 Hz, α = 1.2) ^57^. At inadequate imaging speeds, these diffusive motions would have been missed, as manifested by the distinct trajectories derived by the images taken at 2 Hz and 100 Hz shown in the MSD plot (**Fig. 4c, III**; 2 Hz vs. 100 Hz). Consequently, the actual instantaneous velocity of vesicles during transport, which is approximately 4 µm/s (**Fig. 4c**, IV, 100 Hz), would have been substantially underestimated (estimated as ∼0.5 µm/s in Fig. 4c, IV at 2 Hz, which is in accordance with a previous report ^44^).

**Figure 4:**
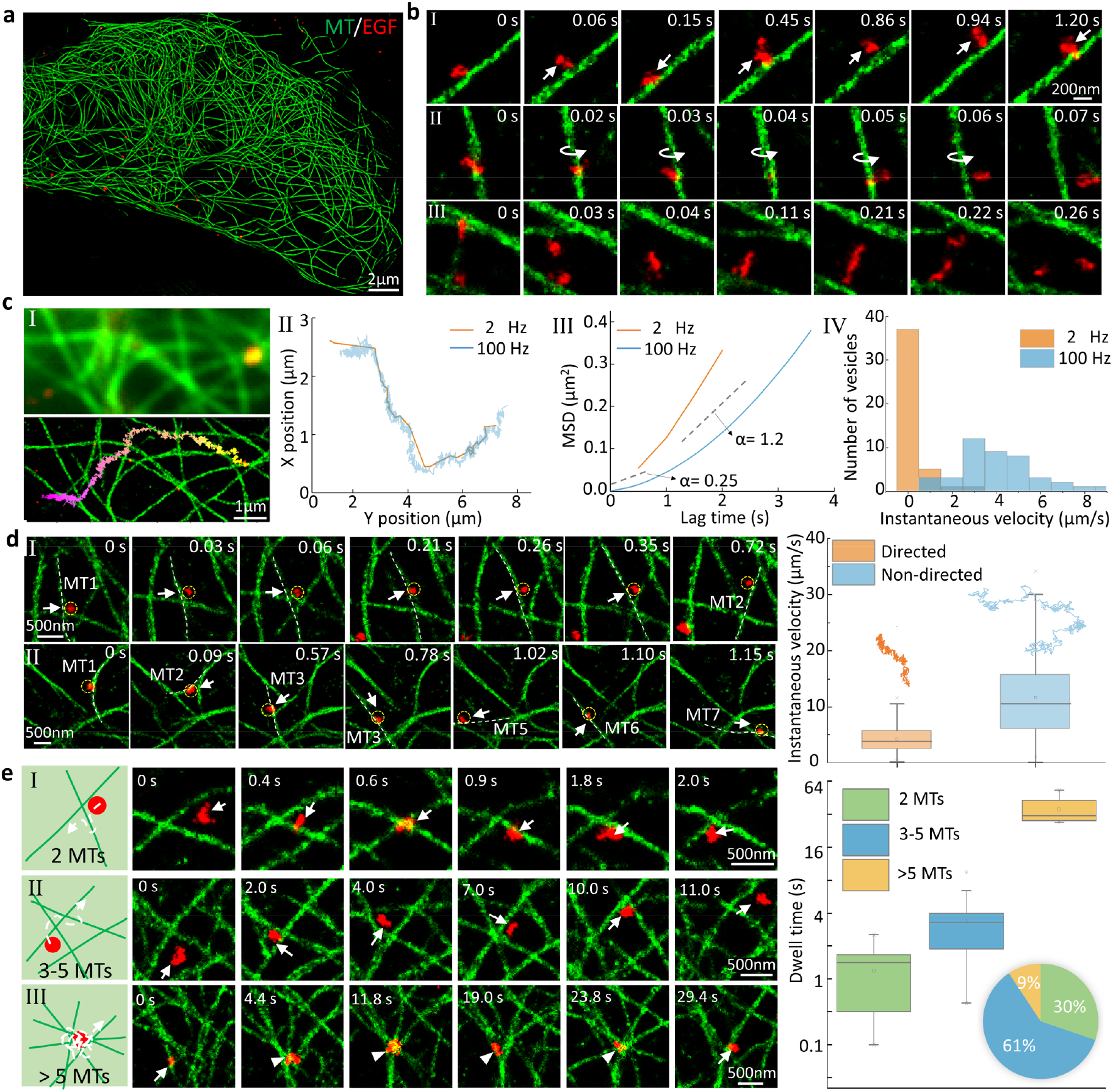
Dual-color real-time SFSRM imaging reveals the microtubule-vesicle interaction. **(a)** Representative dual-channel image of microtubules (green) and vesicles (red) from cells expressing mEmerald-ensconsin and endocytosed QDots655-streptavidin labeled Epidermal Growth Factor (EGF) protein. **(b)** Examples of vesicle transport dynamics: (I) moving back-and-forth along a microtubule; (II) rotating around a microtubule; (III) colliding with another vesicle. **(c)** Example of diffusive motions of vesicle along MT at millisecond scale. (I) Comparison of the dual-color LR and SR image. (II) Comparison of trajectories recorded at 2 Hz and 100 Hz. (III) Mean squared displacement (MSD) analysis of trajectories in (II). MSD reflects the mean-squared-distance ⟨Δr^2^ (τ)⟩ of the vesicle traveled in a certain lag time τ, which typically follows the power-law trend ⟨Δr^2^ (τ) ⟩∝τ^α^, where α indicates the characteristic of the motion. The smaller α is, the more random or diffusive the motion is; while the larger α is, the more directed the motion is. (IV) Statistical comparison of vesicle instantaneous velocity recorded at 2 Hz and 100 Hz respectively. **(d)** Examples of microtubule dynamics resulting in non-directed transport of vesicles. (I) Transverse movement of a microtubule delivers the vesicle to a nearby microtubule. (II) Random fluctuation of surrounding microtubules facilitates the vesicle to switch to different microtubules. The box-and-whisker plot on the right shows the statistical comparison of the instantaneous velocity of the directed/non-directed transport and their representative trajectories. **(e)** Vesicle transport dynamics at different types of microtubule intersections. Based on the number of microtubules involved, the intersections are classified into three categories that are composed of 2 microtubules (I), 3-5 microtubules (II), and more than 5 microtubules (III). The plot on the right shows the statistical comparison of the dwell time at different types of intersections and their percentages in the cell (pie chart).

In addition to these subtle diffusive motions, we also observed nondirected transport, which has been reported in previous studies using single-particle tracking but not fully explained ^44, 61^. Benefitting from our dual-color SFSRM imaging, we observed some MT dynamics that might contribute to nondirected vesicle transport. For example, the transverse movement of an MT can transfer the vesicles attached to it to a nearby MT (**Fig. 4d**, I), and the fluctuations in the surrounding MTs can cause vesicles to switch among different MTs (**Fig. 4d**, II), resulting in nondirected transport (**Supplementary Video 2**, part III). We compared the instantaneous velocities of directed transport and nondirected transport in **Fig. 4d** and found that nondirected movements have an approximately two-fold higher average instantaneous velocity and a four-fold broader range of distribution than directed movements, indicating that these displacements are likely related to MT fluctuations rather than motor-driven movement.

Since MTs are densely distributed, aside from providing tracks for vesicles, they also form intersections which may interrupt vesicle transport. Previous studies have shown that vesicles may pass, pause, switch or reverse at an intersection ^46, 49^. In our experiments, we noticed that all vesicles eventually passed the studied intersections; however, the dwell time varied greatly and largely depended on the complexity of the intersection. Correspondingly, we classified the intersections into three groups based on the number of MTs at each intersection (**Fig. 4e**). For the simplest intersections of two MTs (**Fig. 4e, I**), the vesicle can easily pass through it by climbing over one MT, usually within two seconds; and the MT vibration is unlikely to interrupt vesicle transport. For intersections with 3–5 MTs (**Fig. 4e, II**), the vesicles tended to interfere with the dynamics of nearby MTs. Thus, if the surrounding MTs fluctuate severely, the vesicles are hindered, and the time needed to pass through these intersections ranged from several to ten seconds. For intersections involving more than 5 MTs tethered together (**Fig. 4e, III**), the vesicles were most likely to be trapped at the intersection until the fluctuations of the surrounding MTs became coordinated and the stellate intersection loosened. However, coordinated fluctuations and intersection loosening are highly uncertain, and such processes may take tens of seconds to minutes (**Supplementary Video 2**, part IV). A statistical without missing a beat comparison of the dwell time at different kinds of intersections is shown in **Fig. 4e** (plot). Generally, the more complex the intersection is, the longer the resulting dwell time. For intersections involving more than 5 MTs, the dwell time could be longer than one minute. Fortunately, this kind of intersection only accounts for approximately 10% of all intersections in a cell, whereas more than half of the intersections consist of only 3–5 MTs (**Fig. 4e**, pie chart).

In summary, dual-color live-cell imaging based on SFSRM provides a clear map of the MT network without missing ultrafast dynamic processes, thus successfully capturing vesicle transport details that are oversimplified in most vesicle transport models in which vesicle transport along MTs is normally described as rapid and directed movement. The diverse MT-vesicle interactions indicate that both the MT topology and dynamics can influence vesicle transport, revealing the underlying MT regulation of complex vesicle transport behaviors, such as diffusive motion and nondirected transport; moreover, the various modes by which vesicles pass MT intersections were observed. Our observation that the dynamics of MTs seem to enhance nondirected transport and interfere with directional transport is in good agreement with the results of a previous study by Giannakakou et al.^62^, who reported that the suppression of MT dynamics enhanced nuclear-targeted cargo P53 accumulation near the cell nucleus.

### SFSRM is robust to different imaging systems and different samples

In the above demonstration, we used a Zeiss Elyra 7 microscope as the live-cell imaging system. Here, we demonstrate that SFSRM can also be applied in different live-cell imaging systems without retraining the network model. We validated the robustness of our network based on a commercial confocal microscope (Confocal sp8, Zeiss). Compared to that used for WF imaging, a confocal microscope requires a longer time (2.5 s/frame) to obtain a dual-channel image due to the point scanning strategy used. Here, we recorded the EGF receptor (EGFR) protein transport dynamics at 0.4 Hz for over 10 minutes (**Supplementary Video 3**, part I). Long-term observation allows us to discover some long-time-scale phenomena. For example, as shown in **Fig. 5a**, the EGFR protein gradually accumulates in endosomes, which appear as ring structures in the image. We noticed that the MTs tended to generate local grids to trap the endosomes, as shown in the zoomed-in view in **Fig. 5a**. Our counting results indicate that nearly two-thirds (136/210) of endosomes are trapped in such local MT grids (**Supplementary Video 3**, part II). These traps will actively participate in the transport (**Fig. 5b**) and fusion processes of the endosomes (**Fig. 5c**), and the morphology of these grids will dynamically change in response to the endosome shape (**Supplementary Video 3**, part II). In addition to confocal microscope, we also demonstrate that SFSRM can be directly applied in other imaging systems, such as TIRF and light-sheet microscopes (**Supplementary Fig. 11**).

**Figure 5:**
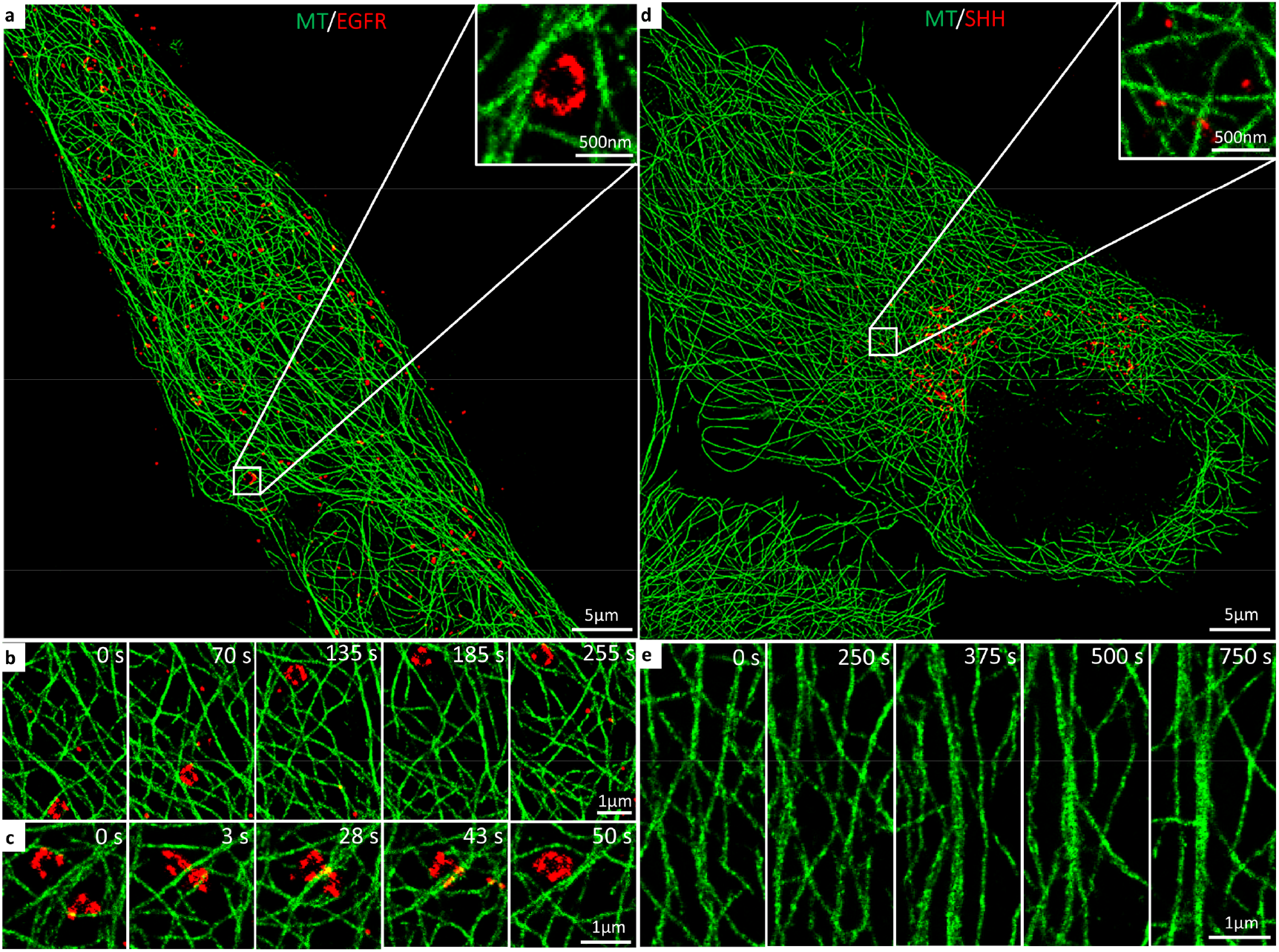
SFSRM cross-system/cross-sample application. **(a)** SFSRM reconstruction of a confocal image of microtubules (green) and EGFR-carrying vesicles (red). Inset: an endosome accumulated with EGFR proteins is trapped in a local microtubule grid. **(b)** Example of the transport process of a trapped endosome. **(c)** Example of the fusion process of trapped endosomes. **(d)** SFSRM reconstruction of a confocal image of microtubules (green) and SHH-carrying vesicles (red). Inset: SHH-carrying vesicles have much smaller size compared with EGFR-carrying vesicles. **(e)** Example of a microtubule bundle generation process.

We further confirmed that SFSRM can be applied to image similar biological structures without retraining the neural networks. The networks trained with EGFR protein images are directly used to reconstruct SR images of sonic hedgehog (SHH) protein (MW: 45 kDa), which is relatively smaller than the EGFR protein (MW: 180 kDa). A video (**Supplementary Video 3**, part III) was taken to study the SHH protein exocytosis process; notably, the SHH protein generated in the perinuclear region of the cell is continuously transported to the cell periphery (**Fig. 5d**). Benefiting from this long-term observation, we were able to capture the entire generation process of a microtubule bundle (**Fig. 5e**). We further investigated the function of the microtubule bundle in vesicle transport. We recorded 18 vesicles transported along this bundle after it was generated. In comparison, only two vesicles traveled through this region before generation (**Supplementary Video 3**, part IV), suggesting that the bundle facilitated vesicle transport. Our observation is consistent with previous results ^63, 64^ and suggests that bundles are generated by acetylated MTs with more kinesin binding sites which aid the vesicle transport.

By implementing SFSRM based on WF and confocal microscopes, both fast and slow temporal dynamics during the vesicle transport process are clearly resolved. The real-time observation of fast dynamics with subtle details significantly enhances our understanding of vesicle transport in live cells, reflecting the capability of SFSRM to facilitate biological research involving live-cell dynamic processes due to the high spatiotemporal resolution of this approach and its adaptivity to different samples and systems.

## Discussion

We have developed and demonstrated an SFSRM method for reconstructing SR images with single-molecule resolution from LR images obtained from conventional microscopes. High-speed single-frame image conversion is achieved by dual subnetworks that progressively improve the image SNR and resolution; enormous resolution and accuracy improvements are achieved by combining an effective network architecture, a multicomponent loss function, and prior information regulation. The adaptation of SFSRM to real-time live-cell imaging pushes the limit of fluorescence microscopes to an unprecedented spatiotemporal resolution with a limited photon budget and circumvents the possible tradeoffs among the spatial resolution, imaging speed, and light dose, thus enabling live-cell imaging at a state-of-the-art resolution, an unprecedented speed, and reduced phototoxicity levels while promoting cross-system modality.

In this work, we implemented SFSRM with a WF microscope for the noninvasive visualization of cargo transport dynamics in a dense MT network, which is challenging for conventional microscopes lacking sufficient spatial or temporal resolutions. The high-frequency transverse vibration of MTs revealed by real-time live-cell imaging via SFSRM can enhance our understanding of the intracellular transport environment. In addition, surprisingly dynamic phenomena during vesicle transport occur within tens of milliseconds, such as diffusive motions along MTs, swinging on MTs, switching between MTs, or diverse behaviors at the MT intersections, can be clearly resolved by the SFSRM. Many of these processes are revealed in relation to the complex topology and high-frequency dynamics of local MTs and have not been seen before. We further confirmed the robustness of SFSRM by directly applying the trained networks to different imaging hardware platforms (such as a confocal microscope) and different biological processes with similar topological structures (such as SHH protein exocytosis). The results demonstrate the potential of SFSRM in investigating other subcellular processes that necessitate interpreting temporal dynamics in the context of ultrastructural information, which may open doors to new discoveries in live-cell imaging involving organelle dynamics ^65^ and interactions ^66^, viral infection ^67^, cellular uptake and the intracellular trafficking of nanoparticles ^61^. Moreover, SFSRM reconciles the conflict between high-resolution and high-throughput microscopy, making it suitable for large-scale, high-workload super-resolution imaging, such as whole-genome imaging ^68^, imaging-based assays of cell heterogeneity ^69^, phenotype profiling ^70^, and drug screening ^71^.

Furthermore, the potential of SFSRM can be further extended to achieve even higher spatiotemporal resolutions. As a learning-based method, the spatial resolution of SFSRM is determined by the SR images it learns from. In our case, all three trained networks have achieved comparable resolutions with the GT images (**Fig. 1, Supplementary Figs. 2** and **8**). Although the highest resolution measured in our experiment was approximately 20 nm (**Supplementary Fig. 8**), this could be further improved if the networks were trained with higher-resolution images, such as those obtained from a MINFLUX microscope ^72^. The temporal resolution that can be achieved in imaging depends on multiple factors, including the probe quantum yield, illumination intensity, the detection efficiency, and even the data transmission speed of the system. In live-cell imaging, the temporal resolution is always sacrificed to achieve an adequate image SNR and observation duration. For instance, we decreased the temporal resolution from 100 Hz to 20 Hz for the long-term whole-cell imaging (**Supplementary Video 2**, part I). Fortunately, with more and more high-quantum-yield labels (e.g., structural modification of fluorescent dyes ^74^ and quantum dots ^75^) available, the temporal resolution could be further improved. In addition to the spatial/temporal resolution improvements, the adaptation of SFSRM to 3D cases ^76, 77^ and/or cases with additional fluorescence channels ^78^ would improve the capability for imaging even more complex biological phenomena.

Nevertheless, the success of SFSRM lies in the use of adequate super-resolution training data. As a data-driven method, the SFSRM approach can learn the statistical patterns of the data and then transform a LR image to a HR image according to the learned mapping. However, this method cannot be used to predict new topological structures that have not been learned before. To adapt to different structures with limited training data, the transfer learning strategy is proven to be capable of accelerating network convergence ^79^. To explore structures beyond the resolution of the current fluorescence super-resolution microscopies, correlative microscopy, which combines fluorescence light microscopy with cryo-electron tomography ^80, 81^, could be further explored to obtain training images with even higher resolutions. Considering these high-cost advanced imaging systems are not ubiquitous in most biological laboratories, we expect, as more public SR image databases become available ^38, 73^, learning-based super-resolution methods will be advanced, and SR imaging can be available to general laboratories.

## Methods

### SFSRM network

The networks of our SFSRM, including the SEN and SRN, are based on the ESRGAN generator ^21^, which includes 23 residual-in-residual dense blocks used to map low-resolution images to super-resolution images. By inheriting the basic architecture of SRGAN^20^, this network performs most computations in the LR feature space, hence reducing complexity and achieving high stability without requiring batch normalization (BN) layers ^21^. The original ESRGAN is designed for a single RGB image. When it was applied to a grayscale image, we found that the network will easily crash at the beginning of or during the training process if we duplicate the grayscale image three times to generate a fake RGB input. Therefore, we adopted a single-channel ESRGAN generator. To incorporate the prior information provided by the edge map, we added another input channel to the network. The input LR image and the corresponding edge map are initially concatenated to generate a two-channel input to the generator. Similarly, duplicated grayscale images are used as fake RGB inputs to the well-trained VGG network ^82^ for feature map extraction.

To generate high-resolution details while maintaining high fidelity, the network is trained with a multicomponent loss function, as follows.

1. Content loss evaluates the L1-norm distance between an estimated SR image G(x_i_) and a ground truth (GT) image y. L1-norm loss focuses on pixel differences, thus allowing the network to quickly converge but often resulting in a blurred image.

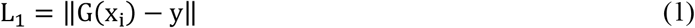

MS-SSIM measures the structural similarity of SR and GT based on luminance, contrast, and structure at different scales. The computation of MS-SSIM is detailed in the assessment metrics section. Here, we focus on the construction of the loss function. MS-SSIM loss is defined as:

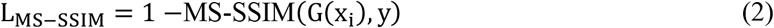

Content loss is a hybrid of MS-SSIM loss and L1-norm loss:

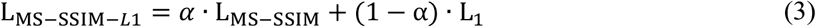

where α is used to balance the contributions of MS-SSIM loss and L1-norm loss and is empirically set as α = 0.84 ^83^.
2. Perceptual loss L_Percep_ is used to measure feature distance differences in the estimated SR image and corresponding GT image. Features are extracted by a VGG network ^82^ pretrained for material recognition and that is good at texture extraction.

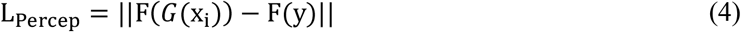

where F represents the feature extraction network.
3. Adversarial loss estimates the probability that the discriminator input x is real or fake. Discriminator D outputs D(x_r_, x_f_) =σ(C(x_r_) − Ex_f_ [C(x_f_)]), where x_r_ and x_f_ are the GT image and generator output, respectively, σ is the sigmoid function, C(x) is the nontransformed discriminator output, and Ex_f_ [·] represents taking the average for all fake data in the minibatch. GAN loss for the generator is formulated as

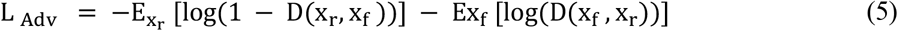
4. Frequency loss compares the frequency distance difference between an estimated SR and the original GT image:

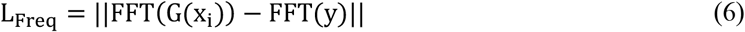

where FFT is the fast Fourier transformation function. We used all frequency components when processing simulation data without noise in the GT image and 75% of frequency components when processing experimental data containing noise in the GT image.

When using the SEN for signal enhancement, the network only uses LR images as single-channel inputs. The training of the SEN includes two steps:

1. Training with MS-SSIM-L1 loss for approximately 100,000 minibatch iterations at a 3×10^−4^ learning rate

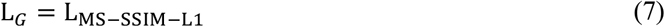
2. Training with MS-SSIM-L1 loss and perceptual loss for 20,000 to 50,000 minibatch iterations at a 1×10^−4^ learning rate.

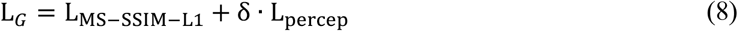

When using the SRN for super-resolution restoration, the network uses both LR images and edge maps as inputs. The training process also includes two stages. The first stage uses the same loss function as the SEN, and the second stage uses the following loss function with a 5×10^−5^ learning rate for approximately 10,000 minibatch iterations.

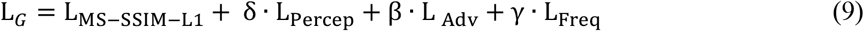

In our experiments, we empirically set δ, β, and γ to δ = 0.1, β = 0.001, and γ = 0.01, respectively.

### Assessment metrics

Multiscale structure similarity (MS-SSIM) ^84^ quantifies the similarity of two images and is an improvement of SSIM ^85^, which assesses the similarity between two images, x and y, based on three factors: luminance l(x, y), contrast c(x, y), and structure s(x, y).

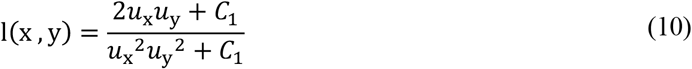

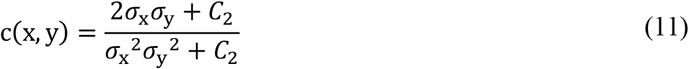

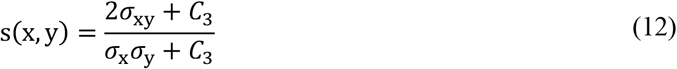

where *C*_1_, *C*_2_ and *C*_3_ are small constants given by *C*_1_ = (*K*_1_*L*)^2^, *C*_2_ = (*K*_2_*L*)^2^, and *C*_3_ = *C*_2_/2. Here, is the dynamic range of pixel values, and *K*_1_ and *K*_2_ are two scalar constants.

The general form of SSIM is defined as:

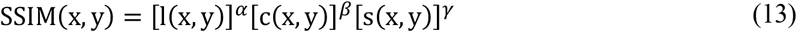

where α, β, and γ are parameters used to define the relative importance of the three components and are set to 1 in most cases ^85^.

MS-SSIM is calculated by iteratively applying low-pass filters, downsampling the filtered image result by a factor M and then calculating the SSIM index of the scaled images. The overall MS-SSIM evaluation is based on combining the measurements at different scales:

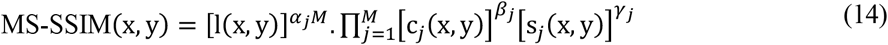

where *α*_*j*_, *β*_*j*_, and *γ*_*j*_ are used to adjust the relative importance of different components ^84^.

The normalized root-mean-square error (NRMSE) quantifies the absolute difference between two images, x and y, and is calculated as follows:

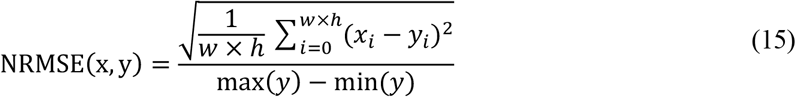

Signal density is computed from a GT image by first conducting binarization for the image to extract the signal-containing pixels and then calculating the ratio of the number of signal-containing pixels to the total number of pixels in the image.

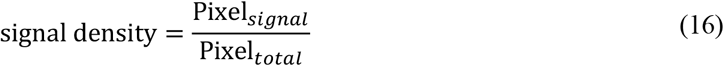

### Image simulation

Simulated polymer chains in a 10 × 10 µm^2^ region were generated in MATLAB. The polymer density was set to 50 polymers per image to mimic a densely distributed MT network. The GT image was created by fitting the fluorophore positions to an image with a pixel size of 10 nm and convolved with a Gaussian kernel with a standard deviation of 1.27 pixels. For the GT image, no noise and a uniform background were used. Similar to the process of generating the GT image, the corresponding LR image was generated by fitting the fluorophore positions to images with a pixel size of 100 nm and then performing convolution with a Gaussian kernel with a standard deviation of 1.27 pixels. In addition to the background, Poisson noise and read noise were added to the LR image.

### Sample preparation

#### Cell culture and transfection

The Beas2B cell line was grown in Dulbecco’s Modified Eagle Medium (DMEM) (Gibco) supplemented with 10% fetal bovine serum (Gibco) and 1% penicillin/streptomycin at 37 °C. The plasmid constructs used in this study included EGFR-mCherry, SHH-RFP, and 3XmEmerald-ensconsin.

#### Staining microtubules in fixed cells

Beas2B cells cultured on coverslips after 24 hours were stained according to the approach in ^86^. Briefly, cells were first washed with cytoskeleton buffer (10 mM MES of pH 6.1, 150 mM NaCl, 5 mM EGTA, 5 mM D-glucose, and 5 mM MgCl2) three times, fixed with 0.6% paraformaldehyde with 0.1% glutaraldehyde and 0.25% Triton, and diluted in CB buffer for 1 minute. Then, 4% paraformaldehyde and 0.2% glutaraldehyde were added, and the resulting mixture was diluted in CB buffer for 15 minutes. After washing three times with 1× PBS, cells were incubated for 10 min in 0.1% NaBH4 to reduce background fluorescence due to glutaraldehyde, and another washing step with PBS was performed. To quench reactive cross-linkers, cells were incubated in 10 mM Tris for 10 min, followed by 2 washes with PBS. Then, cells were permeabilized in 5% BSA and 0.05% Triton X-100, diluted in PBS for 15 min and then incubated with 1:500 mouse anti-α-tubulin antibody (Sigma, T6199) for 1 hour, followed by three washes with PBS. Cells were then incubated with 1:500 Alexa Fluor 647 goat anti-mouse IgG (Invitrogen A-21236) for 1 hour. Finally, the cells were washed with PBS three times and preserved in a standard photoswitching buffer that contained 50 mM Tris of pH 7.5, 10 mM NaCl, 0.5 mg/mL glucose oxidase, 40 μg/mL catalase, 10% (w/v) glucose, and 1% (v/v) β-mercaptoethanol for single-molecule imaging.

### Experimental data acquisition

#### Data acquisition from fixed cells

The experimental training data for fixed cells were obtained from a home-built super-resolution localization microscope ^87^ based on an inverted microscope (Nikon Ti Eclipse) equipped with a 100 × 1.49 NA TIRF objective (Nikon Apo TIRF). Excitation was provided by a 500 mW 656 nm laser (CNI, MRL-N-656.5–5500 mW), and images were acquired by EMCCD (Andor, IXon-Ultra) with a 16 μm pixel size. When performing single-molecule imaging, a 1.5× telescope was used, resulting in a 106 nm effective pixel size. For training data acquisition, a WF image of every field of view was first acquired at low illuminance, and then the laser intensity was increased to the maximum to obtain single-molecule images. For super-resolution imaging, an optimal focus system and a home-built drift-correction system were used to correct system drift. The software was provided by NanoBioImaging Ltd. The frame rate was set to 30 frames per second, and 20,000 frames were acquired per super-resolution image.

#### Data acquisition from live cells

The live-cell data were acquired from different systems, and the image effective pixel size was adjusted to approximately 100 nm. Specifically, the data shown in **Figs. 3** and **4** were acquired from a commercial Zeiss Elyra 7 microscope in HILO mode with a 60×/1.46 oil objective. For an FOV size of 25.6 ×25.6 µm^2^, we recorded dual-color live cell images at 100 Hz with 15 W/cm^2^ illuminance for 5000 time points, and for whole-cell imaging with an FOV size of 60×50 µm^2^, due to the data transmission limitation of the system, we used a 20 Hz imaging speed for 5000 time point recordings; in this process, the illumination intensity was reduced to 3 W/cm^2^. The data in **Fig. 5** were acquired with a Zeiss SP8 confocal microscope at 3 W/cm^2^ illuminance with a 63×/1.4 oil objective. We recorded 300 time points at 0.4 Hz for an FOV of 51.2 ×51.2 µm^2^.

#### Image processing

The single-molecule image sequences were analyzed with the ThunderSTORM ^88^ plug-in in FIJI. The super-resolution reconstructed images were obtained at 5× magnification for MT images and 10× magnification for vesicle images. To generate the training data, the LR images were processed by a custom code to extract the edge map. To generate the training pairs of LR images, edge maps, and GT images, the LR images and edge maps were interpolated at a scale of 1.25× based on bicubic interpolation. The intensity of all images was normalized to the range of 0–255. Then, the images were split into small blocks of size 256×256 to correspond to the size of the GT images (64×64 for LR images and edge maps). Finally, approximately 1000 training pairs were used to train the network for the simulated polymer images; approximately 300 training pairs were used to train the network for the experimental microtubule images; and approximately 600 training pairs were used to train the network for the experimental vesicle images.

#### Online content

Any methods, additional references, and supplementary information are available on line.

## Supporting information

The real-time imaging of microtubules in a live cell via SFSRM shows the diverse microtubule dynamics.

The dual-color real-time imaging of EGF protein and microtubules in live cells via SFSRM reveals the vesicle-microtubule interaction dynamics.

The demonstration of applying SFSRM to different microscopes/samples.

## Acknowledgements

We thank Dong Li (University of Chinese Academy of Sciences) for providing 3XmEmerald-ensconsin plasmid. We also thank HKUST Bioscience Central Research Facility and Super-Resolution Imaging Center for essential equipment support. We gratefully acknowledge financial support from the Research Grants Council of Hong Kong under the General Research Fund (GRF; 16205818 and 16205619 to S.Y.; and 16102921, 16102218, 16103319, and 16104020 to Y.G.).

## Author contributions

S.Y. and S.D. conceived the project. S.Y., S.D., and Y.G. supervised the research. S.Y., S.D., and R.C. designed the experiments. X.T., T.L., Y.S., J.W., B.C. prepared samples. Z.S. and R.C. performed experiments. R.C. analyzed the data with conceptual advice from S.Y., S.D. and Y.G. R.C. wrote the manuscript with input from all authors under the supervision of S.Y, S.D. and Y.G. All authors discussed the results and commented on the manuscript.

