## Supplementary Note and Figures for "Deep-Learning Super-Resolution Microscopy Reveals Nanometer-Scale Intracellular Dynamics at the Millisecond Temporal Resolution"

### Content

**Supplementary Note 1:** Signal-to-noise ratio (SNR) estimation and measurement

**Supplementary Figure 1:** Subpixel edge map of a microscopic image based on the fluorophore's radial symmetry

**Supplementary Figure 2:** Validation of super-resolution network (SRN) on an experimental microtubule image acquired from a fixed cell

**Supplementary Figure 3:** Validation of super-resolution network (SRN) on different signal density levels

**Supplementary Figure 4:** Evaluation of SFSRM on different signal-to-noise ratio (SNR) levels

**Supplementary Figure 5:** Evaluation of SFSRM on different imaging systems

**Supplementary Figure 6:** Comparison of SFSRM with representative methods on public dataset

**Supplementary Figure 7:** Evaluation of SFSRM on different spectra

**Supplementary Figure 8:** Representative super-resolution (SR) reconstructions of EGF and SHH images by SFSRM

**Supplementary Figure 9:** System drift and microtubule transverse vibration

**Supplementary Figure 10:** Examples of microtubule morphology changes due to vesicle transport

**Supplementary Figure 11:** Representative reconstruction results of SFSRM from images obtained with common imaging systems

**Supplementary Video 1.** The real-time imaging of microtubules in a live cell via SFSRM shows the diverse microtubule dynamics such as bend, growth or shrink, and transverse vibration. Beas2B cells were transfected with mEmerald-ensconsin plasmid. A region of 25.6 ×25.6 $\mu m^2$ was chosen to record the video at 100 Hz for 5000 time points. The illumination intensity was set as 15 $W/cm^2$ during imaging.

**Supplementary Video 2.** The dual-color real-time imaging of EGF protein and microtubules in live cells via SFSRM reveals the vesicle-microtubule interaction dynamics during the transport process including diffusive motions, nondirected transport, and passing through microtubule intersections. Beas2B cells were transfected with mEmerald-ensconsin plasmid and then incubated with Qdot655-labeled EGF protein to allow EGF endocytosis. For the whole cell imaging in the first part of the video, a region of 60×50 $\mu m^2$ was chosen to record the video at 20 Hz for 5000 time points. The illumination intensity was set as 3 $W/cm^2$ during imaging. While for the high-speed dual-color imaging, a region of 25.6 ×25.6 $\mu m^2$ was chosen to record the video at 100 Hz for 5000 time points. The corresponding illumination intensity was set as 15 $W/cm^2$.

**Supplementary Video 3.** The demonstration of applying SFSRM to different microscopes/samples. The first two parts of the video are the long-term dual-color imaging of EGFR protein and microtubules in live cells, from which we observe that microtubules form grids and participate in endosome transport. The second two parts of the video are the dual-color imaging of SHH protein and microtubules in live cells, which captures the microtubule bundle generation process. For both experiments, the video of a region of 51.2 ×51.2 $\mu m^2$ was recorded for 300 time points at 0.4 Hz. The illumination intensity was set as 3 $W/cm^2$.

**Supplementary Note 1: Signal-to-noise ratio (SNR) estimation and measurement**

**Theoretical estimation of SNR for live-cell images**

Here we discuss how our experimental conditions including the exposure time (t), illumination density (P), and fluorescent probes for which we mainly consider its quantum yield (Φ) and duty cycle (η), will affect the acquired image signal-to-noise ratio (SNR) in live-cell experiments. Generally, the photons emitted by fluorescent proteins or probes, after passing through the optical system, are captured by a detector (e.g., a camera). For a single fluorescent molecule, its emission photon is determined by the absorbed photons $Q_{ab}$ and quantum yield Φ of the detector:

$$Q_{em} = Q_{ab} \cdot \Phi \quad (1)$$

where $Q_{ab}$ is proportional to the amount of the photons $Q_I$ arrived at the sample plane and determined by the absorbance A and the on-off duty cycle η of the fluorescent molecule.

$$Q_{ab} = \eta(1 - 10^{-A})Q_I \quad (2)$$

When A is very close to zero which can be normally satisfied since the absorbance of a molecule is very small, the equation above can be reformulated as:

$$Q_{ab} = (\ln 10)\eta Q_I A \quad (3)$$

According to the Beer-Lambert law, the absorbance A of a molecule can be described by its molar attenuation coefficient ε as:

$$A = \varepsilon c L \quad (4)$$

where c is the molar concentration of the fluorescent molecule and L is the pathlength. Here we consider L as the depth of focus (DOF) of the objective:

$$L = \frac{\lambda}{2NA^2} \quad (5)$$

where $\lambda$ is the excitation wavelength and NA is the numerical aperture of the objective.
Hence $Q_{ab}$ can be denoted by

$$Q_{ab} = (\ln 10)\eta\varepsilon c L Q_I \quad (5)$$

Given the illumination density P, region of interest S, exposure time t, and photon energy of the excitation laser hυ , $Q_I$ can be further denoted as

$$Q_I = PSt/h\upsilon \quad (6)$$

Taken together, the emission photon number can be calculated as follows:

$$Q_{em} = (\ln 10)PSt\eta\varepsilon cL\Phi/h\upsilon \quad (7)$$

If the collected photons, further go through the dichroic mirror and bandpass filter of the detection path with a transmission coefficient $T_D$ and $T_B$, The emission photons collected by the objective with a capture efficiency $\gamma = \frac{1}{2}(1 - \sqrt{1 - (NA/n)^2})$ [1], where n is the refractive index of the immersion medium, then the detected photon number $Q_{det}$ can be denoted by:

$$Q_{det} = \gamma Q_{em} T_D T_B \quad (8)$$

Due to the inherent shot noise which is generated by the photon variation, and readout noise, dark current noise resulted from the internal process of the camera when converting photon to current, the SNR of the generated can be denoted as:

$$\mathrm{SNR} = \frac{QE.\mathrm{Q_{det}}}{\sqrt{QE.\mathrm{Q_{det}}+N_r+I_d}} \tag{9}$$

where $QE$ is the quantum efficiency; $I_d$ is the dark current noise; $N_r$ is the readout noise.

Here we give an example of using mEmerald as the fluorescent probe and PCO.edge 4.2 sCMOS camera as the detector for calculating the emission photon number and the relative SNR. The parameters used are listed below:

Table 1. The parameters and the corresponding values used in the SNR estimation.

| **Parameter** | **Value** | **Definition** |
|---|---|---|
| Φ | 0.68 | Quantum yield of fluorophore |
| η | 1/3 | Duty cycle of fluorophore [$T_{on}/(T_{on}+T_{off})$] |
| ε | $57500\ cm^{-1}M^{-1}$ | Fluorophore attenuation coefficient |
| c | 10 μM | Fluorophore concentration |
| L | 0.1 μm | Depth of focus of the objective |
| S | $0.1 \times 0.1$ μm$^2$ | Sample area corresponding to one pixel on the detector |
| hυ | $4.4 \times 10^{-19}$ J | Photon energy [$\lambda_{ex}$ =488nm] |
| P | $3\ W/cm^2$ | Illumination density |
| T | 10 ms | Exposure time |
| γ | 0.37 | Capture efficiency of the objective |
| $NA$ | 1.46 | Numerical aperture of the objective |
| $n$ | 1.51 | Refractive index of the immersion medium |
| $T_D$ | 0.9 | Transmission coefficient of the dichroic |
| $T_B$ | 0.95 | Transmission coefficient of the bandpass filter |
| $QE$ | 0.82 | Quantum efficiency of the detector |
| $I_d$ | 0.3 e- | Dark current noise |
| $N_r$ | 0.9 e- | Readout noise |

The detected photon can be calculated following equation (8):

$$Q_{\mathrm{det}} = \frac{\ln 10 \times 3\frac{W}{cm^2} \times 10\mathrm{ms} \times (0.1\mu\mathrm{m} \times 0.1\mu\mathrm{m}) \times \frac{1}{3} \times 57500\frac{L}{mol.cm} \times \frac{10\mathrm{umol}}{\mathrm{L}} \times 0.1\mu\mathrm{m} \times 0.68 \times 0.37 \times 0.9 \times 0.95}{4.4 \times 10^{-19}\ \mathrm{J}} = 9.69$$

$$\mathrm{SNR} = \frac{0.82 \times 9.69}{\sqrt{0.82 \times 9.69 + 0.3 + 0.9}} = 2.63$$

We further estimate the photon emission of three fluorescent proteins used in our experiments listed as follows:

Table 2. Typical parameters of three probes and the estimated SNRs

| | $\lambda_{ex}$ (nm) | $\lambda_{em}$ (nm) | $\Phi$ | ε (cm−1 M−1) | $Q_{\mathrm{det}}$ | SNR |
|---|---|---|---|---|---|---|
| mEmerald[2] | 482 | 509 | 0.68 | 57500 | 9.69 | 2.63 |
| mCherry[3] | 587 | 610 | 0.22 | 72000 | 4.78 | 1.71 |
| RFP[3] | 555 | 584 | 0.41 | 98000 | 11.46 | 2.89 |

The experimental conditions can be adjusted to get a certain image quality for different experiment goals. For example, for long-term observation, we maintained a low illumination intensity of 3 $W/cm^2$ to reduce photobleaching while increasing the exposure time to 50 ms to meet the SNR requirement; while for ultrafast imaging, we increased the illumination intensity to 15 $W/cm^2$ to allow for ultrahigh temporal resolution. However the hardware limitation, for instance, the data transfer speed, as well as photodamage[4] shall also be taken into consideration in order to optimize the photon budget.

**Experimental measurement of SNR for acquired images**

In our work, we use SNR as a metric for quantifying the signal level of the image, which can be obtained as follows:

a. The input image Im is blurred with a Gaussian kernel with a standard deviation of five pixels to generate $\mathrm{Im}_{\mathrm{blur}}$.
b. The signal mask $S_{\mathrm{mask}}$ is extracted by a manually-set threshold of 0.2; and the background mask $B_{\mathrm{mask}}$ is extracted by a manually-set threshold of 0.1.
c. The signal region $S_{\mathrm{pixel}}$ and background region $B_{\mathrm{pixel}}$ of the input image Im are extracted by applying the signal mask $S_{\mathrm{mask}}$ and background mask $S_{\mathrm{mask}}$ corresponding to Im;
d. The signal $\mu$ is obtained by subtracting the averaged signal $S$ by the camera offset $offset$, estimated by calculating the average of the background pixels, while the noise $\delta$ is composed of the signal variance $\sqrt{S - offset}$ and noise variance N, which is N the standard deviation of the background pixels.
e. The SNR of Im is computed according to the following equation:

$$\mathrm{SNR} = \frac{\mu}{\delta} = \frac{S - offset}{\sqrt{S - offset + N^2}} \tag{10}$$

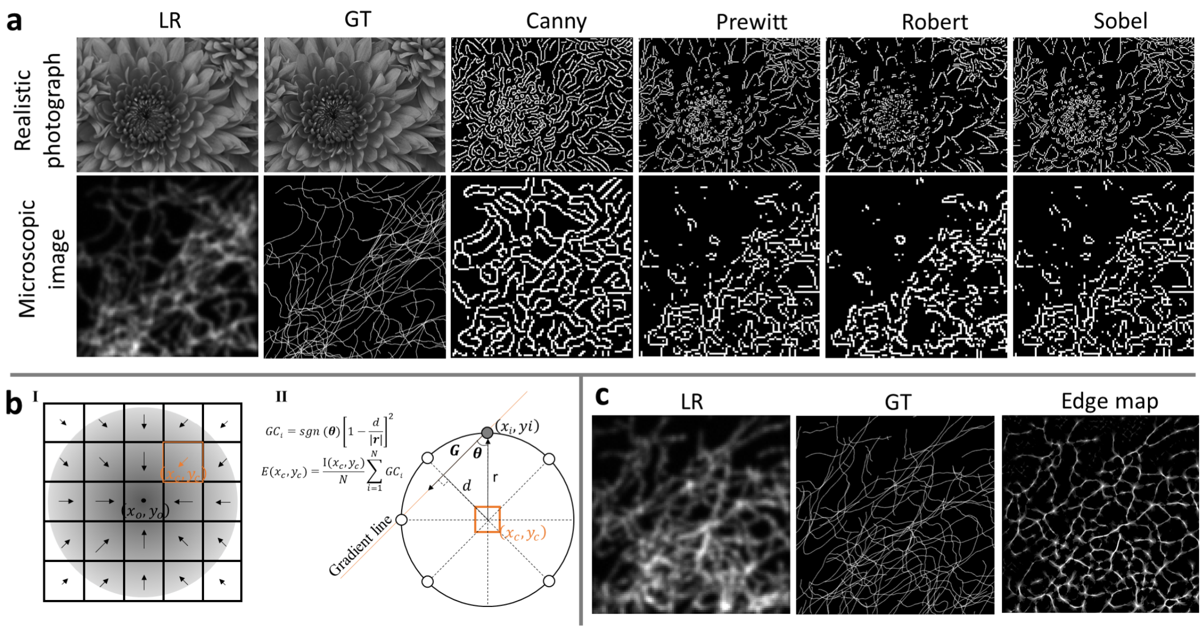


**Supplementary Figure 1. Subpixel edge map of a microscopic image based on the fluorophore's radial symmetry.** **a**, Comparison of the low-resolution (LR) image, ground-truth (GT) image, and edge maps extracted by Canny, Prewitt, Roberts, and Sobel edge detection operator from the realistic photograph and microscopic images. These common edge detectors work well for the realistic photograph however cannot indicate real edges of the microscopic GT image. **b**, Illustration of the calculation of the edge map based on the fluorophore's radial symmetry. (I) The gradient map of a fluorophore located in $(x_o, y_o)$. Since the gradient map is symmetrically distributed and converges to the location of the fluorophore, the local gradient convergence can be utilized to measure the fluorophore signal distribution. (II) For an arbitrary coordinate $(x_c, y_c)$, the edge map value $E(x_c, y_c)$ is determined by the gradient convergence ($GC_i$) index of N coordinates $(x_i, y_i)$ evenly distributed surrounding it with a distance of $r$, where $GC_i$ indicates the angle between the gradient vector $\boldsymbol{G}$ and radical vector **r**. The final edge map $E(x_c, y_c)$ is further weighted by the local fluorescent signal intensity. **c**, An example of the edge map extracted from a microscopic LR image by our subpixel edge detector, which fits well with the true edges in the GT image.

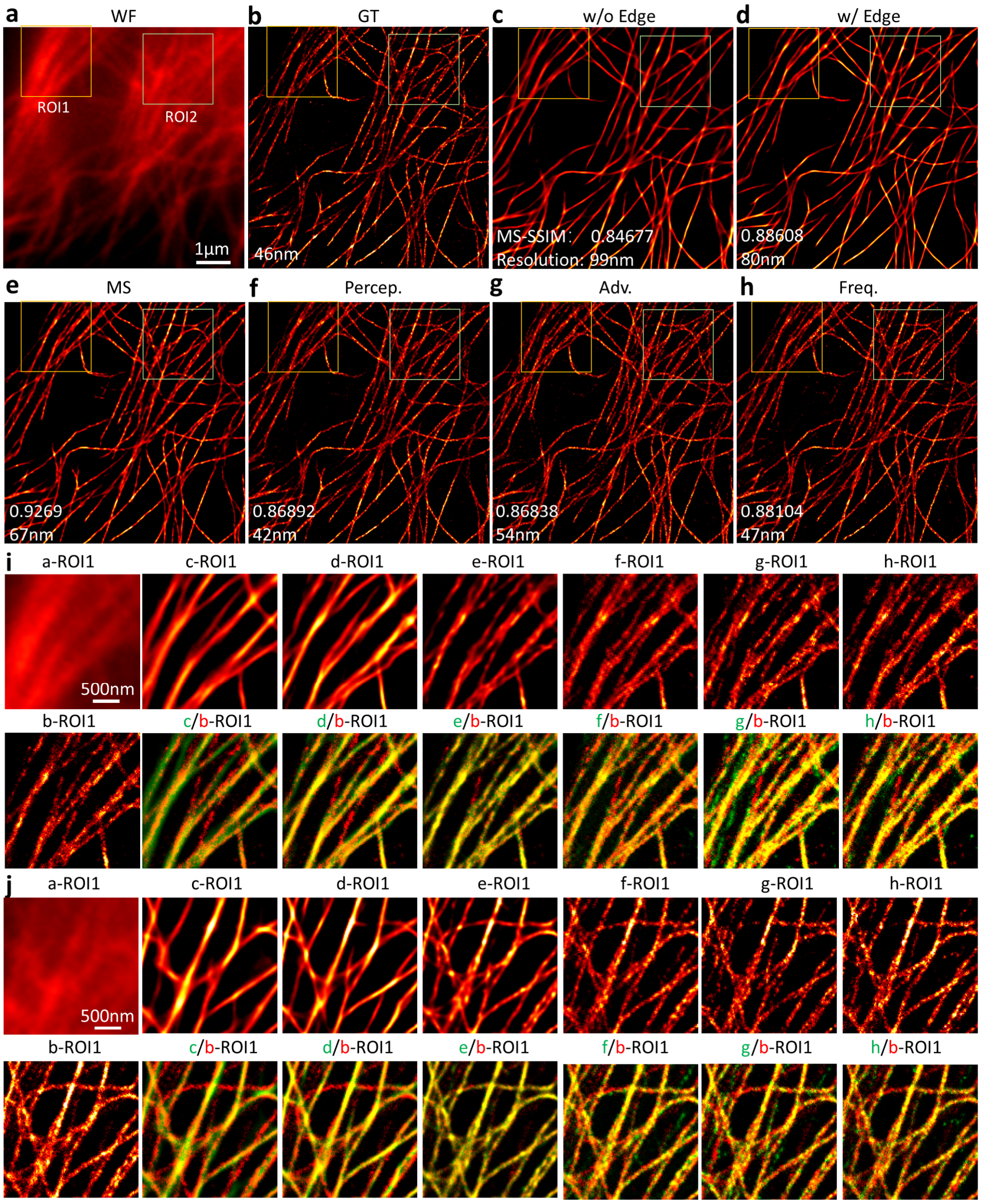


**Supplementary Figure 2**. **Validation of super-resolution network (SRN) on an experimental microtubule image acquired from a fixed cell.** **a**, The WF image of Beas2B cell immunostained by Alexa Fluor 647. **b**, The STORM reconstruction from 20,000 frames of single-molecule images of the same region with **a,** regarded as the GT image. **c-d,** Comparison of networks without (w/o) and with (w/) the edge map as a network input. **e-h**, Comparison between different loss components including MS-SSIM-L1 loss (MS), perceptual loss (Percep.), adversarial loss (Adv.), and frequency loss (Freq.). The reconstruction fidelity of **b-g** is assessed by the multi-scale structure similarity (MS-SSIM) with respect to the GT image and the reconstruction resolution is measured by image decorrelation analysis[1]. **i-j**, Zoom-in views of ROI1 and ROI2 in **a-h** and the merged images of reconstruction results **c-h** (in green) with the GT image **b (**in red).

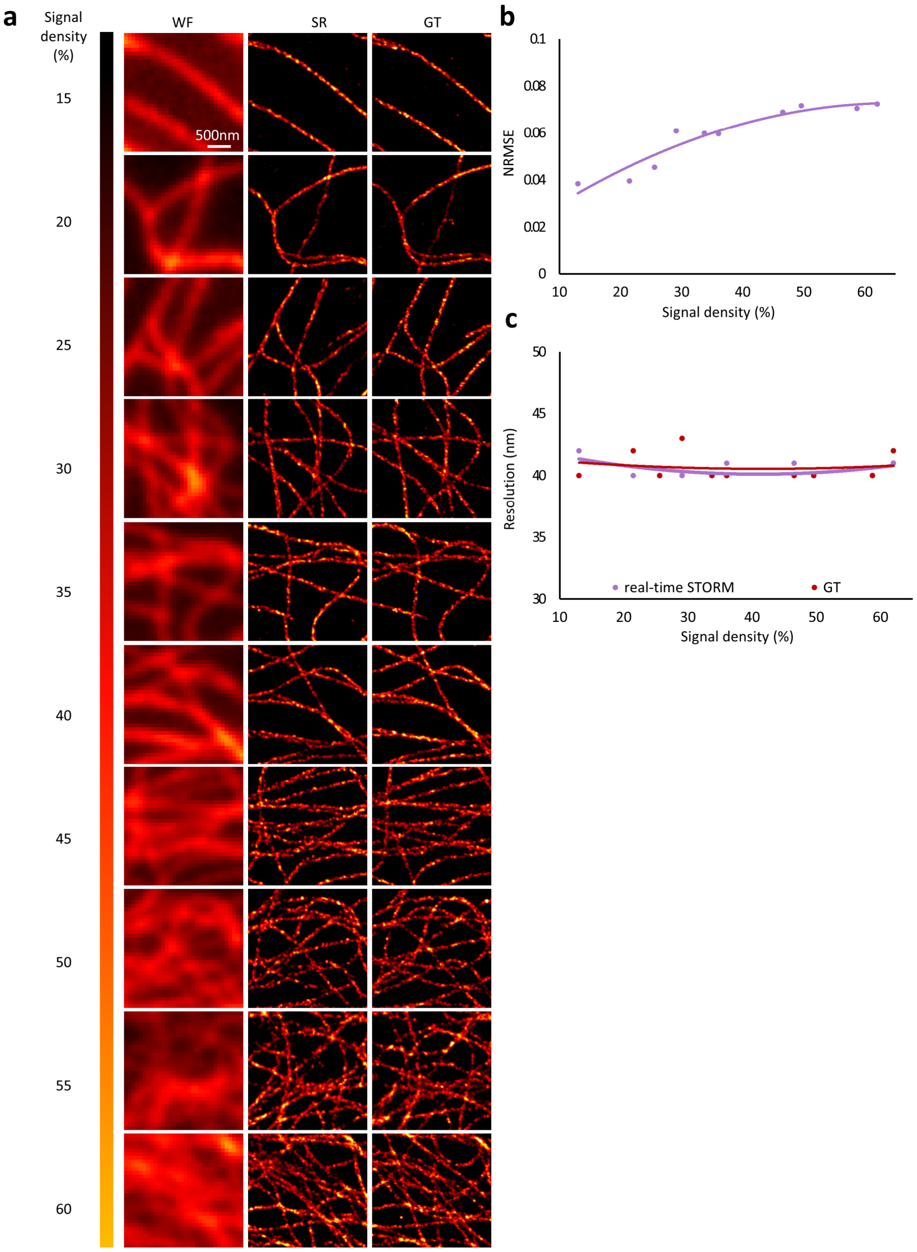


**Supplementary Figure 3: Validation of super-resolution network (SRN) on different signal density levels.** **a**, The first column shows the representative WF images of different signal density; the second column shows network reconstruction noted as SR images; the third column shows the STORM reconstruction from 20,000 frames of single-molecule images of the same region regarded as the GT image. The signal density is calculated by first applying image binarization to the GT image to extract the structure map and then calculating the ratio of the signal-containing pixels to the total pixels of the whole image. **b**, Reconstruction error is quantified by normalized root-mean-square error (NRMSE) of SR images reconstructed from inputs with different signal intensity. **c**, Reconstruction resolution (measured by decorrelation analysis) of SR images reconstructed from inputs with different signal density (purple) and the resolution of corresponding GT image (red).

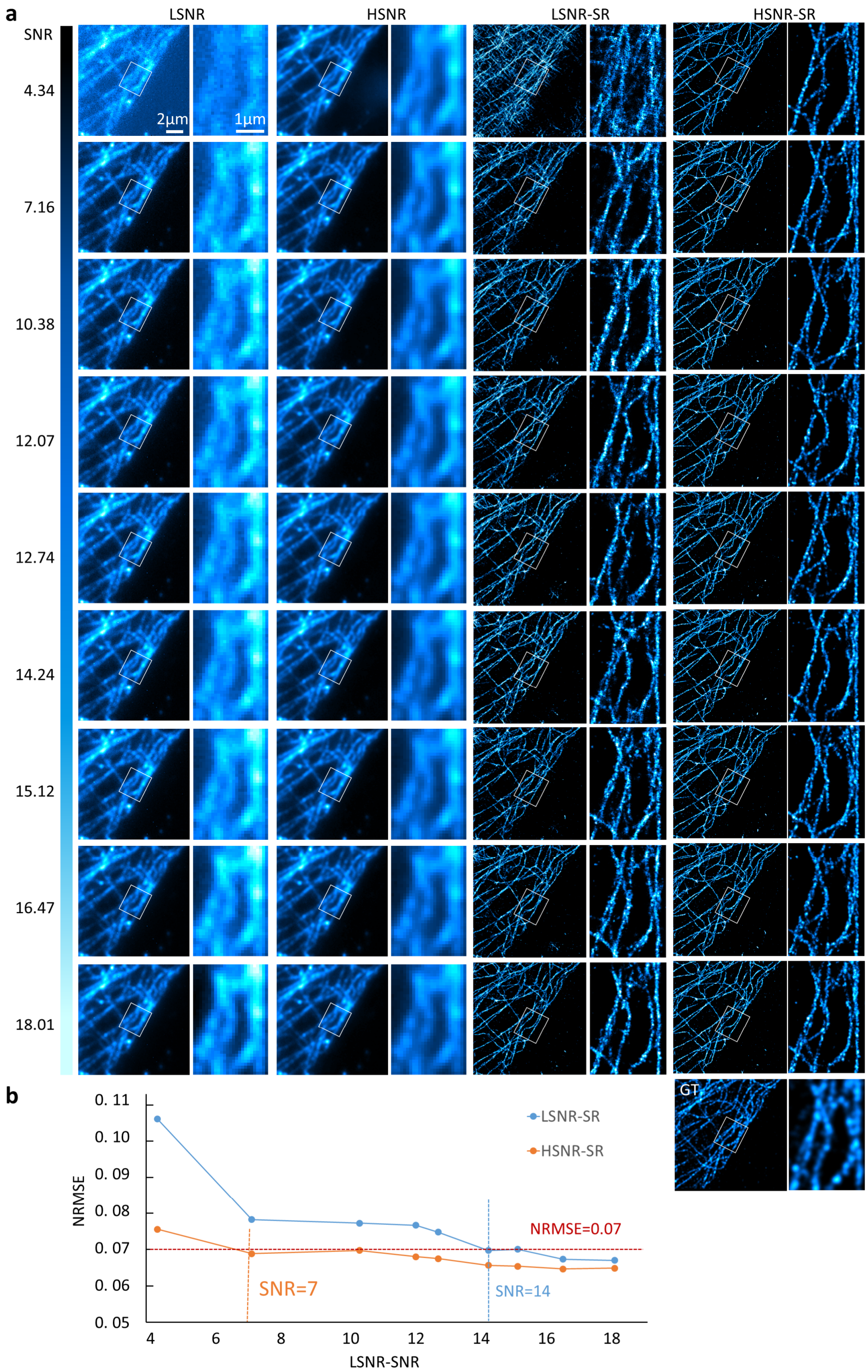


**Supplementary Figure 4**. **Evaluation of SFSRM on different signal-to-noise ratio (SNR) levels. a**, The first column shows the original low SNR image noted as LSNR; the second column shows the intermediate result from the signal enhance network (SEN), noted as high-SNR (HSNR) image; the third column shows the super resolution network (SRN) reconstruction results from the original LSNR images, noted as LSNR-SR; the last column shows the SRN reconstruction results from the HSNR images, noted as LSNR-SR. The last row shows the STORM image of the same region reconstructed from 20,000 frames of single-molecule images, regarded as GT image. **b**, The normalized root-mean-square error (NRMSE) of the SR images reconstructed from the LSNR image (noted as LSNR-SR) and HSNR image (HSNR-SR). Generally, the HSNR-SR image has lower NRMSE compared with the corresponding LSNR-SR. We chose the NRMSE of 0.07 as the cutoff value for getting an acceptable reconstruction fidelity. When the input LSNR image has SNR >14, the NRMSE of the LSNR-SR is lower than the 0.07. In contrast, the NRMSE of HSNR-SR is lower than 0.07 when the corresponding LSNR input SNR > 7. Thus, without SEN, the SFSRM has a minimum SNR request of SNR>14; with SEN, this SNR threshold could be further decrease to 7.

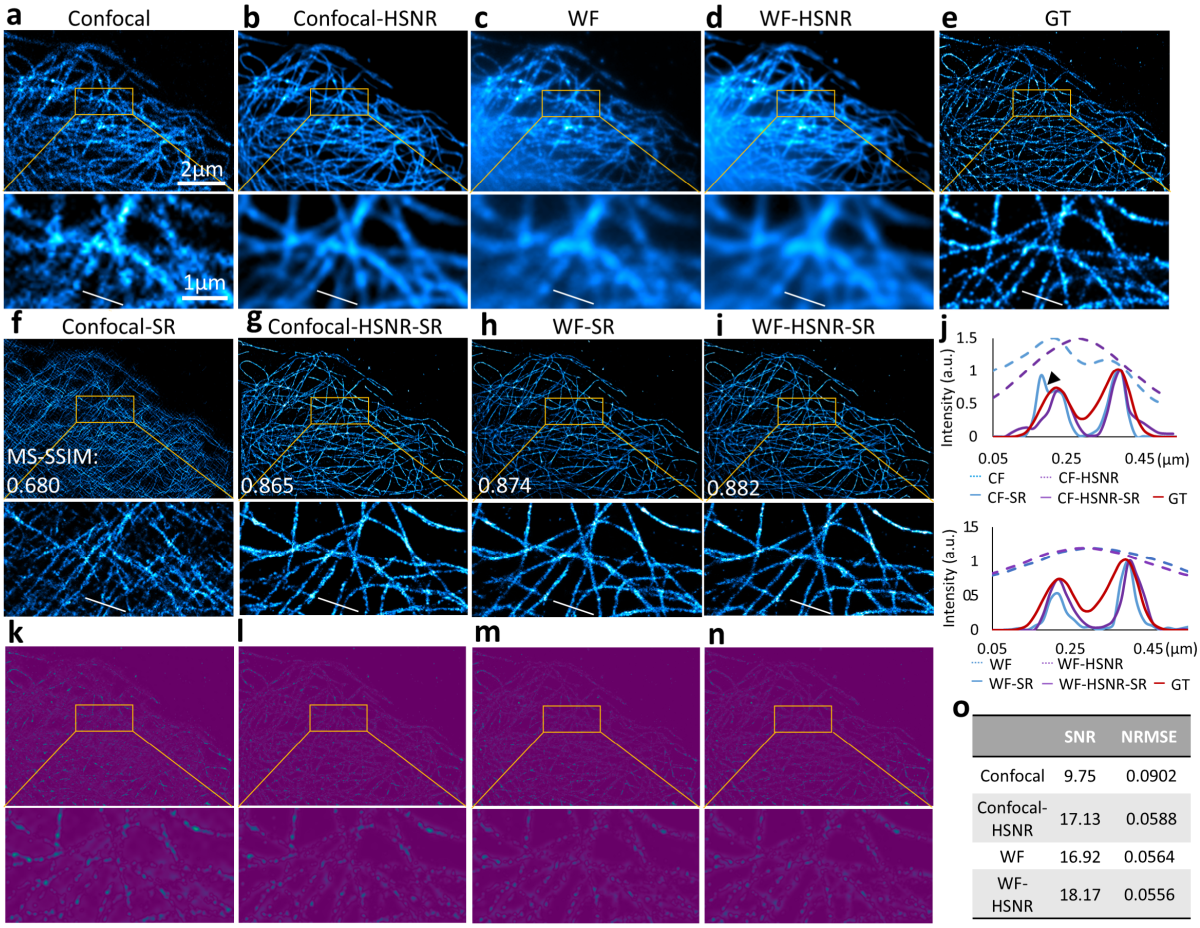


| | SNR | NRMSE |
|---|---|---|
| Confocal | 9.75 | 0.0902 |
| Confocal-HSNR | 17.13 | 0.0588 |
| WF | 16.92 | 0.0564 |
| WF-HSNR | 18.17 | 0.0556 |

**Supplementary Figure 5: Evaluation of SFSRM on different imaging systems. a**, The raw image of microtubules in Beas2B cells immunostained by Alexa Fluor 647 obtained from a Zeiss sp8 confocal microscope. **b**, The HSNR image reconstructed by the signal enhance network (SEN) from the confocal image a. **c**, The raw image of the same sample acquired from a Zeiss Elyra 7 microscope using the highly inclined and laminated optical sheet (HILO) illumination mode, noted as the WF image. **d**, The HSNR image reconstructed by SEN from the WF image. **e**, The STORM image of the same sample, reconstructed from 20,000 frames of single-molecule images, regarded as the GT image. **f-i**, super-resolution (SR) reconstructions of panels **a-d** by the SRN, the reconstruction fidelity of **a-d** is assessed by the multi-scale structure similarity (MS-SSIM) index with respect to the GT image. **j**, Intensity profiles along the white lines in **a-i**. Black arrow in **j** indicates the artifact in **f**. **k-n**, Difference images of panels **f-i** with respect to the GT image. **(o)** Comparison of the normalized root-mean-square error (NRMSE) of the reconstructions in panels **f-i**.

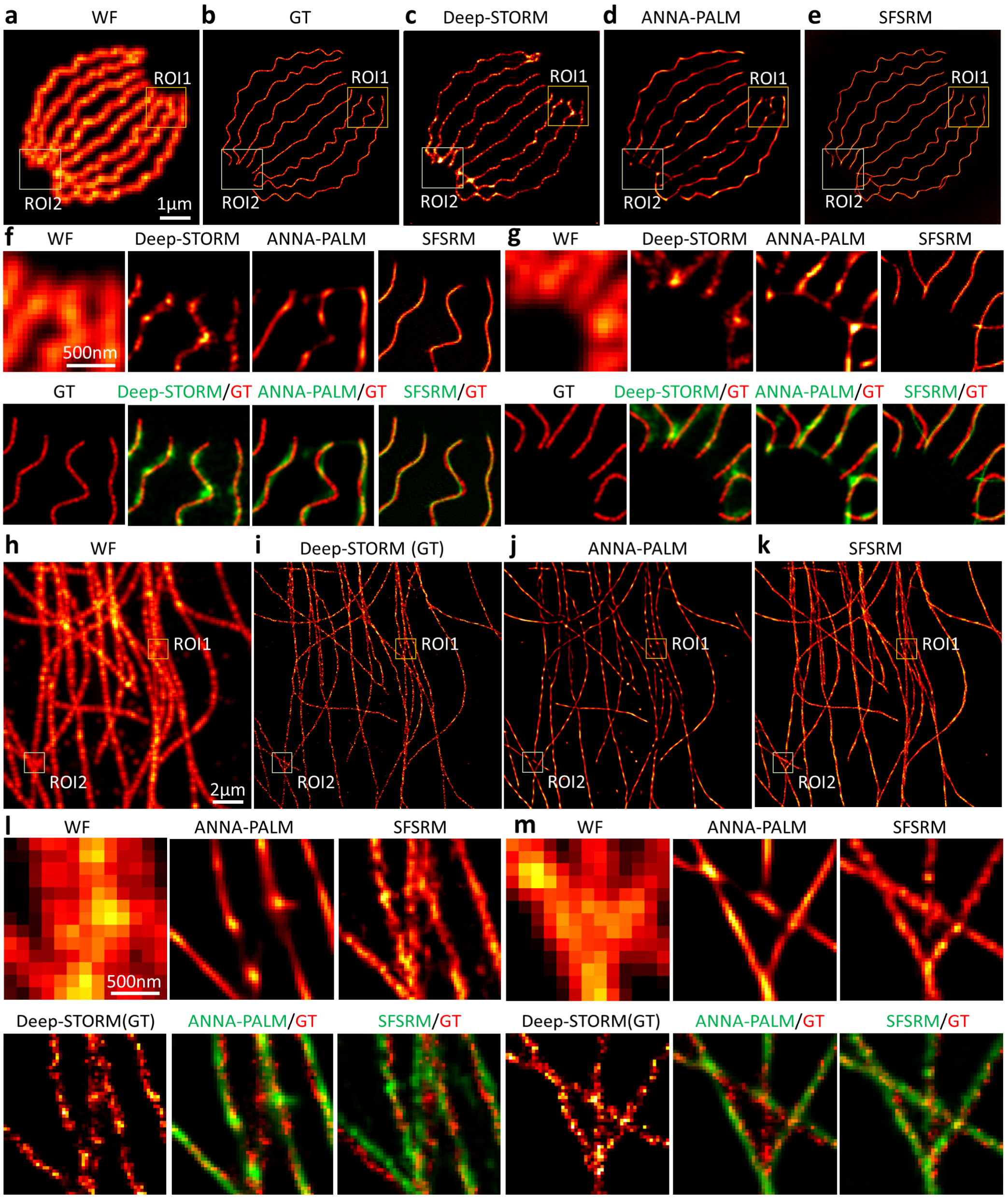


**Supplementary Figure 6: Comparison of SFSRM with representative methods on public dataset. a**, Simulated WF image of microtubules generated by averaging the acquisition stack. **b**, GT image. **c**, Reconstruction by Deep-STORM from 300 frames of densely-distributed single-molecule images. **d**, Reconstruction by ANNA-PALM from the WF image. **e**, Reconstruction by SFSRM from the WF image **a**. **f-g**, Zoom-in views of ROI1 and ROI2 in **a-e** and the merged images of the reconstructions **c-e** (in green) with the GT image **b** (in red). **h**, Experimental WF image of microtubules generated by averaging the acquisition stack. **i**, Reconstruction by Deep-STORM from 300 frames of densely-distributed single-molecule images regarded as the GT image. **j**, Reconstruction by ANNA-PALM from the WF image. **k**, Reconstruction by SFSRM from the WF image. **l-m**, Zoom-in views of ROI1 and ROI2 in **h-k** and the merged images of the reconstructions **j-k** (in green) with GT image **i** (in red).

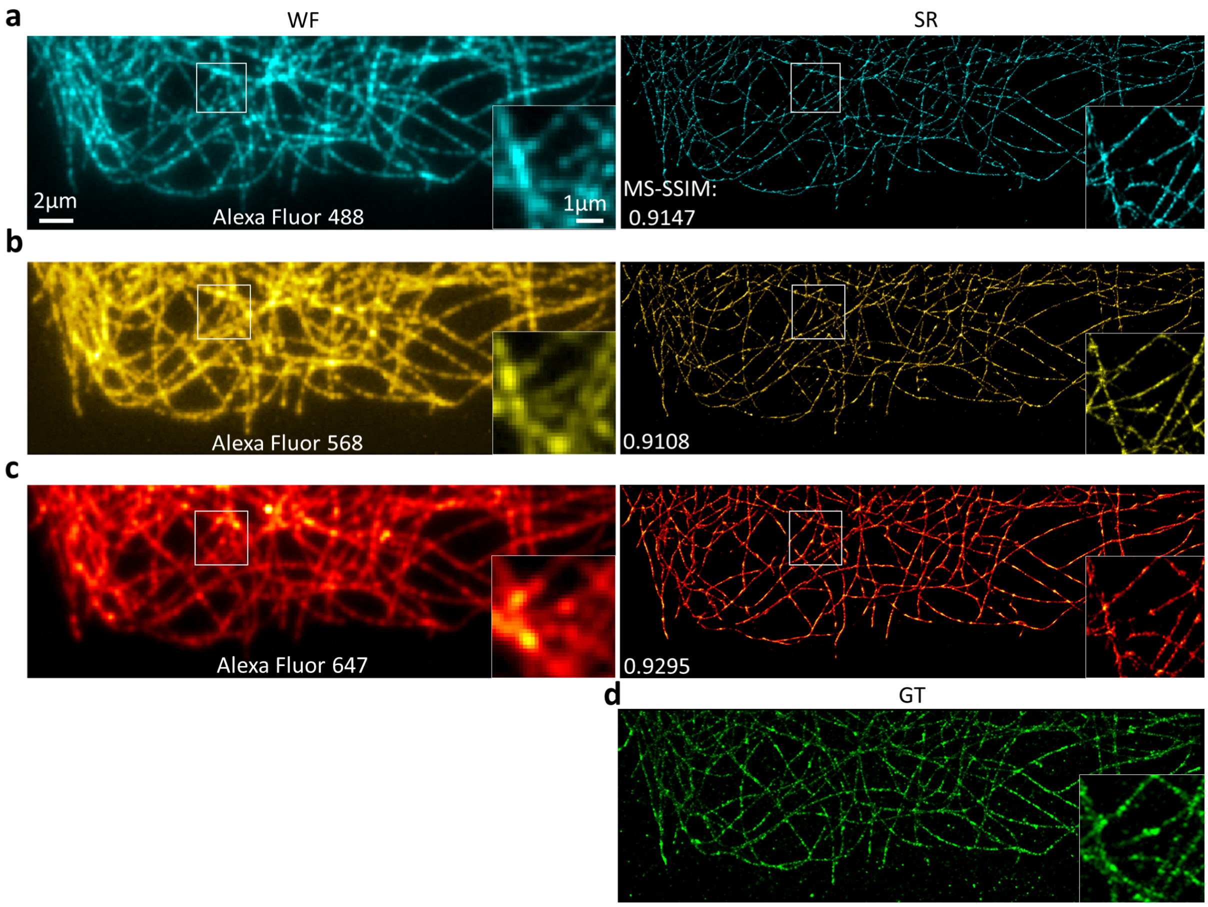


**Supplementary Figure 7: Evaluation of SFSRM on different spectra. a-c**, The WF images (left panel) and SFSRM reconstructions (right panel) of microtubules immunostained by Alexa Fluor 488,568, and 647 simultaneously. The insets show the zoom-in view of the regions highlighted in white box. The reconstruction fidelity assessed by multi-scale structure similarity (MS-SSIM) of the SR images with respect to the GT image. **d**, The STORM image reconstructed from 20,000 frames of single-molecule images, regarded as the GT image.

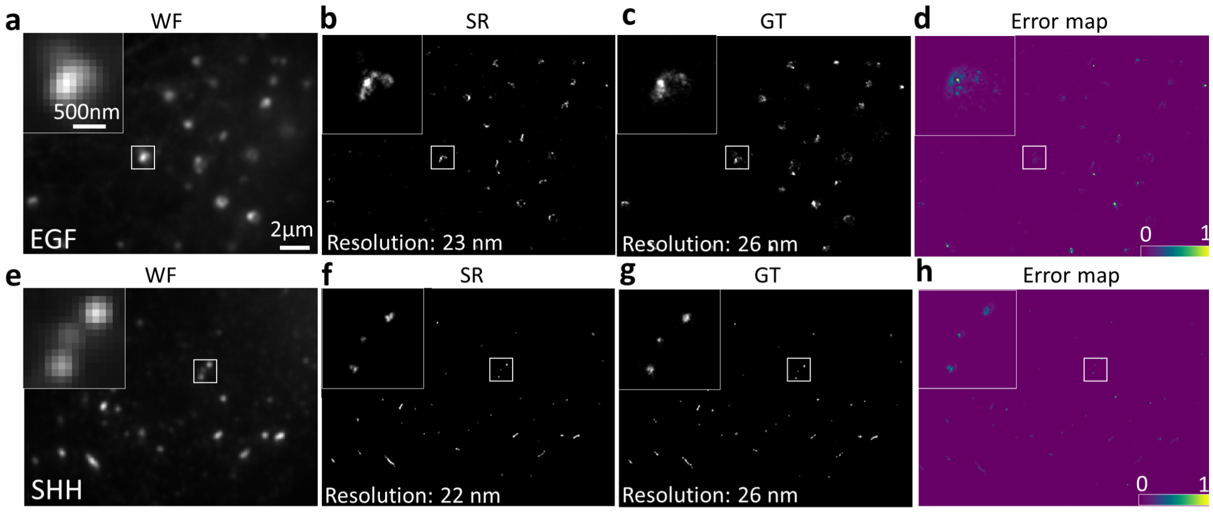


**Supplementary Figure 8: Representative super-resolution (SR) reconstructions of EGF and SHH images by SFSRM. a**, The WF image of vesicles carrying Epidermal Growth Factor (EGF) protein immunostained by Alexa Fluor 647. The inset shows the zoom-in view of the region highlighted in white box. **b**, The SFSRM reconstructed SR image from **a**. Resolution is measured by image decorrelation analysis. **c**, The STORM reconstruction from 20,000 frames of single-molecule images of the same region with **a,** regarded as the GT image. **d**, Difference image of the SR image with the GT image. **e**, The WF image of vesicles carrying Sonic hedgehog (SHH) protein immunostained by Alexa Fluor 647. **f**, The SFSRM reconstructed SR image from **e**. The SFSRM network trained using EGF images is directly used to reconstruct the SHH image. **g**, The STORM reconstruction from 20,000 frames of single-molecule images of the same region with **e,** regarded as the GT image. **h**, Difference image of the SR image with the GT image.

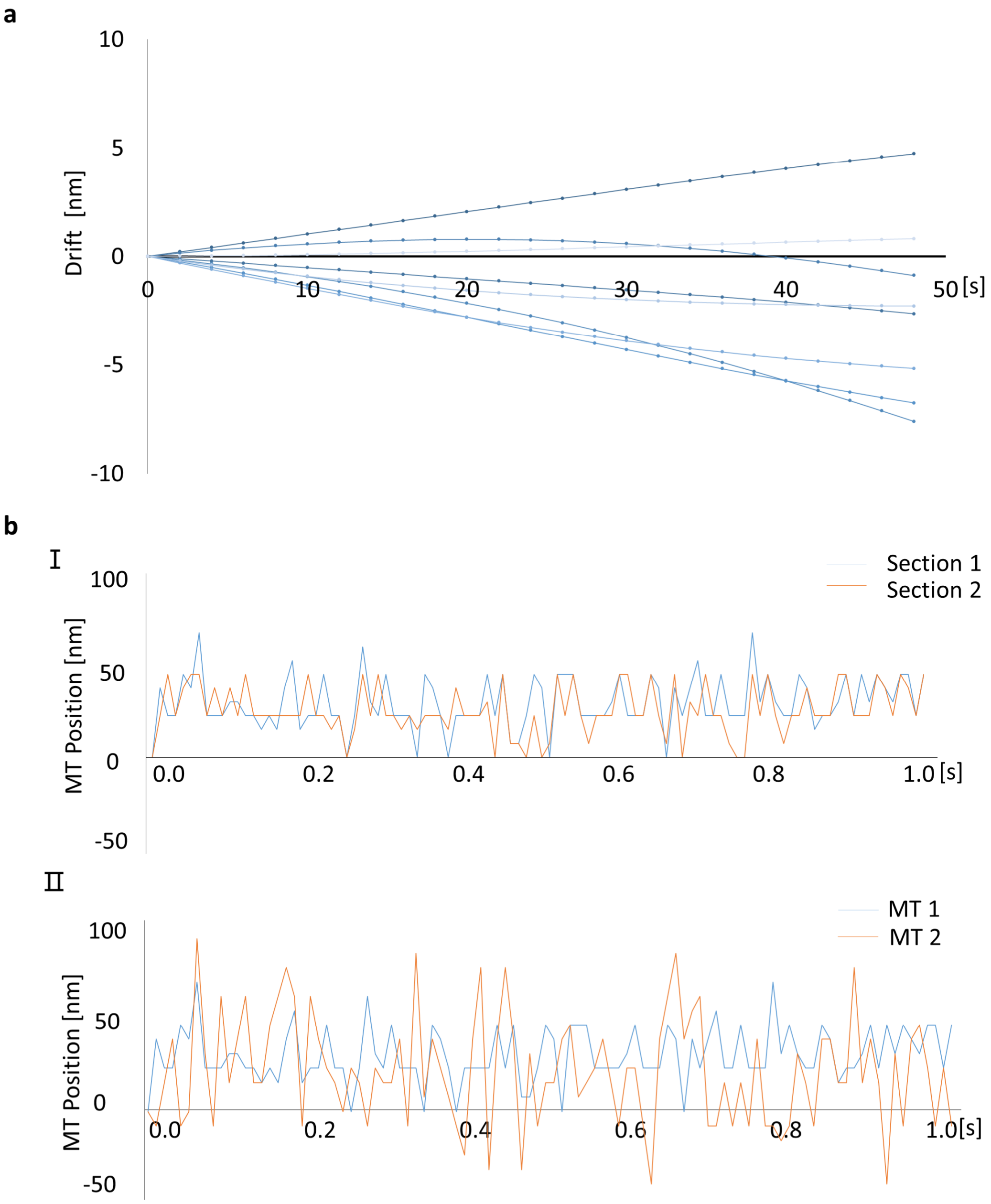


**Supplementary Figure 9: System drift and microtubule transverse vibrations. a**, The system drift in 50 seconds tested using fluorescent beads coated on a coverslip. The experiment was repeated for 8 times, showing that the drift was consistently smaller than 10 nm in 50 seconds, which was the duration of our high-speed live-cell imaging experiment. **b**, (I) The vibrations of two microtubule sections cut from the same microtubule. (II) The vibrations of two different microtubules in the same duration. The cross-correlation of the two curves is 0.91 in (I) and 0.56 in (II). The low cross-correlation of the vibration curves of different microtubules indicates that this vibration is not likely to be the uniform system drift, while the high cross-correlation of the vibration curves of two sections of the same microtubule further confirms this kind of vibration is originated from active motions of an individual microtubule.

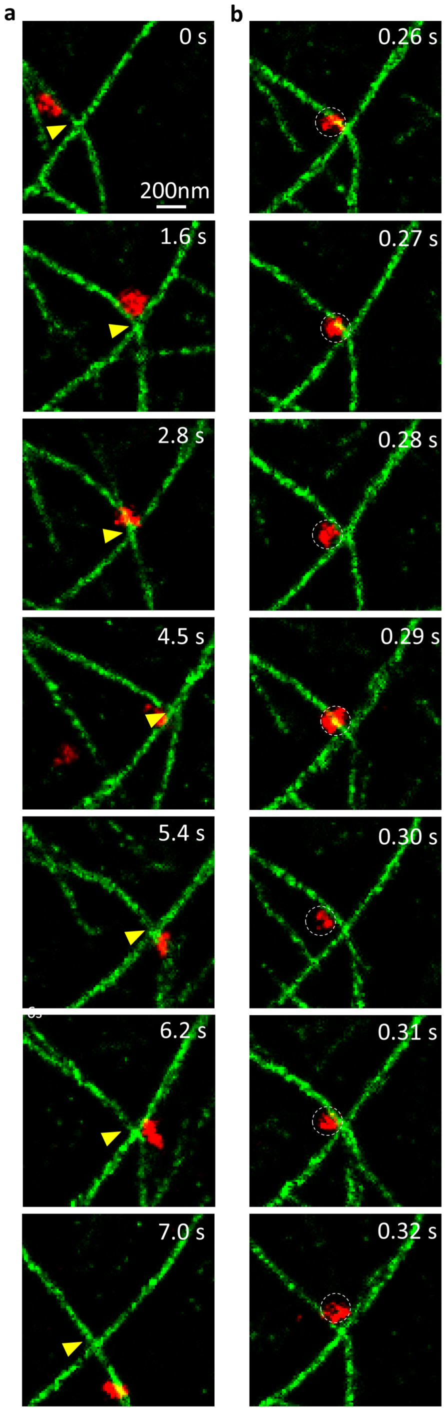


**Supplementary Figure 10: Examples of microtubule morphology changes due to the vesicle transport. a**, The time-lapse images showing that a microtubule bends to its limit positionand returns to the original position. **b**, The time-lapse images showing the unstable vesicle movement.

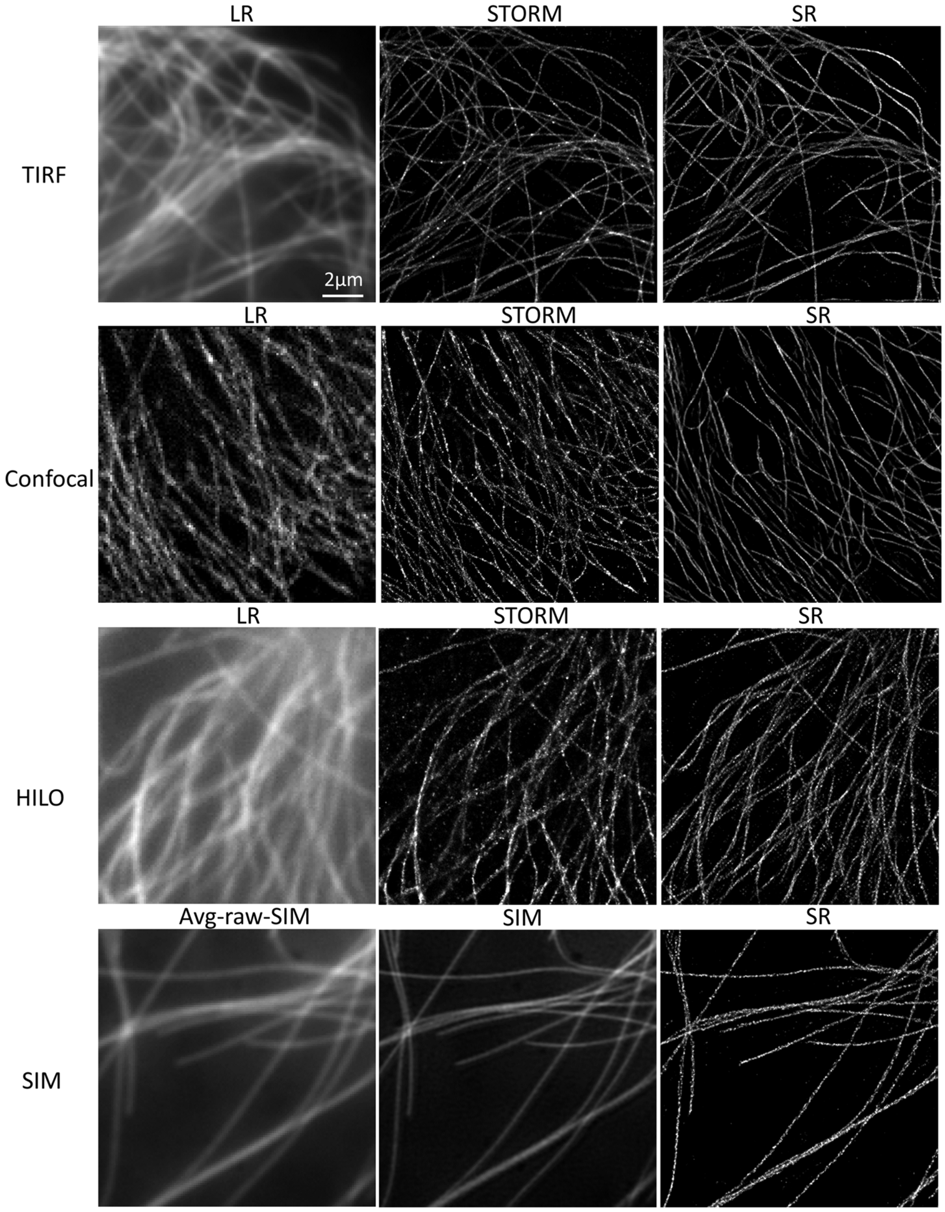


**Supplementary Figure 11: Representative reconstruction results of SFSRM from images obtained with common imaging systems.** The first row shows the raw image of microtubules obtained from a Nikon Ti2 TIRF microscope using Beas2B cells immunostained by Alexa Fluor 647, the corresponding STORM image reconstructed from 20,000 frames of single-molecule images, and the SFSRM reconstruction result from the raw TIRF image. The second row shows the raw image acquired from a Zeiss confocal sp8 microscope, the corresponding STORM image, and the SFSRM reconstruction result from the raw confocal image. The third row shows the raw image acquired from a Zeiss Elyra 7 microscope using HILO mode, the corresponding STORM image, and the SFSRM reconstruction result from the raw HILO image. The fourth row shows the average-intensity-projection image of a raw SIM image provided by Qiao et al.[2], the corresponding SIM reconstruction image also provided by Qiao et al. [2], and the SFSRM reconstruction result from the average-intensity-projection image.
